# CanID: a robust and accurate RNAseq Expression-based diagnostic classification scheme for pediatric malignancies

**DOI:** 10.1101/2025.08.20.671349

**Authors:** Daniel K. Putnam, Alexander M. Gout, Delaram Rahbarinia, Meiling Jin, David Finkelstein, Xiaotu Ma, Jinghui Zhang, David A. Wheeler, Larissa V. Furtado, Xiang Chen

## Abstract

Cancer subtype classification is critical for precision therapy and there is a growing trend of augmenting histopathology testing procedures with omics-based machine learning classifiers. However, analytical challenges remain for pediatric cancer on the scope and precision of the current classifiers as well as the evolving subtype standardization. To address these challenges, we built Cancer Identification or CanID, a stacked ensemble machine learning classification scheme, using the transcriptomic features derived from gene-level RNA sequencing count data as the sole input. CanID was developed primarily from 3203 pediatric cancer samples of 13 solid tumor subtypes and 38 hematologic malignancy subtypes with subtype labels curated without the use of RNA-seq data. The accuracies of independent testing in three independent or external data sets for Solid Tumor and Hematologic Malignancy are 99% and 92–93%, respectively. Notably, CanID was able to classify subtypes challenging for clinical histology evaluation and was robust to both biological and technical challenges, including differences in data collection protocols, class imbalance, potential mislabeled training samples and classes unobserved in training. The high accuracy, robustness, biological interpretability of this transcriptome-based classification scheme represents a valuable approach to advance tumor diagnosis and clinically meaningful stratification of tumor types. CanID can be accessed on GitHub at https://github.com/chenlab-sj/CanID.

## Introduction

Despite advances in the therapy and patient care, pediatric cancer remains the leading cause of death among children with over 16,000 new cases each year in the United States[1], and 400,000 globally[2]. Subtype classification provides critical input for risk stratification which determines treatment intensity as well as eligibility for targeted therapy [3, 4] with the ultimate goal of developing more effective and less toxic therapies for pediatric cancer patients. Traditionally, this has been carried through multiple tests using biomarker-based molecular diagnosis and histopathological approaches. These methods are constrained by the effort required for assay development and validation, as well as the need for iterative testing. As genome-wide profiling has become a standard practice in pediatric oncology testing, subtype classification is increasingly performed by applying validated computational analysis to genome-wide omics data, rather than designing assays for individual biomarkers. For example, fusion gene expression[5] and multi-gene expression signature analyses[6, 7] have been incorporated into routine clinical diagnosis for specific cancers. Moreover, DNA methylome-based classifiers have been developed for accurate tumor subtyping in brain tumors[8] as well as early tumor detection in cell-free DNA[9].

The high accuracy achieved by methylation-based classifiers which were developed for analyzing pediatric central nervous system (CNS) tumors using the random forest algorithm has established the feasibility of deriving an unbiased machine learning-based subtype classification scheme. A limitation of the DNA methylation platform is the lack of direct functional interpretability of the underlying signal measured by methylation probes. By contrast, features extracted from gene-based whole transcriptome sequencing (i.e. RNA-seq) analyses have the potential to generate a biologically interpretable classification scheme. RNA-seq is also a key genomic data type generated by major pediatric cancer genomic research initiatives [e.g. St Jude/Washington University Pediatric Cancer Genome Project (PCGP[10]) and the National Cancer Institute (NCI) Therapeutically Applicable Research to Generate Effective Treatments (TARGET[11]] as well as clinical sequencing programs (e.g. Genome4Kids[12], INFORM[13], ZERO[14]). This enables the assembly of a sufficient number of RNA-seq samples from the public databases, which is essential for training and validating classifiers for very rare pediatric cancer subtypes. When combined with methods for visualizing high dimensional data, such as t-distributed stochastic neighbor embedding (tSNE), RNA-seq has proven useful for classifying tumors in research settings[15–17]. Leveraging RNA-seq data in this way is particularly useful for discerning tumor subtypes without relying on known biomarkers.

Machine learning approaches are well suited for classification tasks, as they are designed to learn a mapping between a set of observations (e.g. input omics features) and an outcome (e.g. cancer subtypes) from an existing reference dataset (training set). If the learned mapping, is generalizable, it can be applied to classify new query samples. Although the classification performance of a machine learning algorithm is typically measured by numerical metrics such as prediction accuracy, precision, recall, F_1_ score, area under the receiver operating characteristic curve (AU-ROC), and confusion matrices, in many real-world applications, numerical metrics alone fail to capture a complete description[18]. In addition to achieving high accuracy, it is often desirable for well-performing machine learning models to reveal how a certain classification (or decision) is made, especially in misclassified cases.

Recently, multiple machine learning classifiers for pediatric cancers using RNA-seq have been developed [19–21], providing subtype classification that is independent of biomarker-based assays and has the potential to streamline molecular diagnosis and narrow the scope of histopathological testing. However, to date, these approaches have either been limited in their scope of B-cell acute lymphoblastic leukemias (B-ALL) cases included [20], restricted to certain classifiable subtypes of B-ALL [21] or applicable only to samples prepared using the polyA-enriched mRNA-seq protocol [19].

Here we report the development of CanID, a unified classification scheme designed to classify pediatric tumors using diverse RNA-seq data. CanID normalizes systematic differences between the major RNA-seq protocols, corrects potential batch effects between sample cohorts, and extracts biologically important features using principal component analysis (PCA). To predict the type for individual tumor transcriptomes, we built a stacked ensemble classifier from five base classifiers. To demonstrate the utility of this approach, we focused on pediatric non-CNS solid tumors (ST) and hematologic malignancies (HM) due to the lack of well-accepted molecular classifiers and we compared its performance with that of a recently published pediatric tumor classifier, OTTER [19]. OTTER’s developers trained the model using an ensemble of convolutional neural networks on whole-transcriptome gene expression data from poly(A) RNA-Seq samples. They identified OTTER classes using a hierarchal clustering algorithm, RACOON.

We built CanID models and classified ∼4800 samples consisting of 13 subtypes of pediatric STs and 38 subtypes of pediatric HMs including acute myeloid leukemia (AML), B-cell acute lymphoblastic leukemia (B-ALL), and T-cell acute lymphoblastic leukemia (T-ALL), profiled by mRNA-seq or total RNA-seq [22]. We compiled these samples from 7 research initiatives, 2 clinical sequencing programs, and 2 external data sources. (Supplementary Tables 1–5). Users can run CanID as a standalone method or integrate it into a workflow that incorporates ancillary biomarker information such as gene fusion or somatic mutations, which they can also compute from RNA-seq.

## Methods

### RNAseq data sets and subtype designation

We obtained all RNA-seq data from public resources including (1) 3202 samples profiled by 7 research studies and 2 clinical sequencing initiatives from St. Jude Cloud (SJCloud) [23] (https://www.stjude.cloud/); (2) NCI’s Therapeutically Applicable Research to Generate Effective Treatments initiative [11] (TARGET; n = 1555; https://ocg.cancer.gov/programs/target/data-matrix); (3) 8 rhabdoid tumor samples from Australian’s The Zero Childhood Cancer program[14] for the purpose of expanding the sample size of rhabdoid tumors used for training and testing; and 4) 61 samples from a clinical pilot study[24]. Details are summarized in Supplementary Tables 1–4.

We obtained subtype information of the SJCloud clinical samples from clinical diagnosis guided by the World Health Organization (WHO) criteria[25], key molecular findings, and input from St. Jude pathologists with domain expertise in the relevant tumor type. For research samples, we derived subtype information primarily from the corresponding publications (Supplementary Table 5, [10, 12, 16, 26–33]). Because both the clinical and research samples were collected over a 10-year period, some diagnostic criteria may have changed. To ensure consistency in diagnostic labeling, we harmonized diagnostic subtype annotations for tumors in this study by re-reviewing all cases, examining clinical notes from patients’ electronic medical records, confirming the presence of diagnostic biomarkers when appropriate, and consulting with pathologists who specialize in hematologic and solid tumors. In some cases, this process led to subtype reclassification. For example, several samples labeled as T-lineage ALL of HOXA (TALLHOXA) subtype in a recent publication [16], was reclassified as acute myeloid leukemia with re-arranged KMT2A (AMLKMT2A) based on a review of the original publication [34] and analysis of their gene expression profiles: SJMLL021, SJMLL022, SJMLL012, SJMLL011, SJMLL013, SJMLL018, SJMLL020. Similarly sample SJMLL030007, also labeled as TALLHOXA in the same publication [16], was labeled as AMLNPM1 based on gene fusion status reported in a different study [24]. Subtype designations for the TARGET cohort were based in part from the Genomic Data Commons (https://portal.gdc.cancer.gov) and from published literature [16, 35].

### RNA data processing

We measured gene expression levels using RNA-seq feature counts. For RNA-seq data hosted on St Jude Cloud, we obtained the corresponding “Feature Count” files from the St Jude Cloud Genomic Platform (https://platform.stjude.cloud/). We computed RNA-seq feature counts for TARGET and ZERO using the same St Jude Cloud pipeline (https://stjudecloud.github.io/rfcs/0001-rnaseq-workflow-v2.0.0.html) [23]. Briefly, we mapped RNA-seq reads to the hg38 genome build (GRCh38_no_alt) with the STAR aligner in two-pass mode[36] and annotated them with GENCODE v31 gene models (https://www.gencodegenes.org/human/release_31.html). We then generated raw gene-level counts with HTSeq-count[37], also using GenCode v31 gene models.

### Filtering Protocol-Sensitive Genes

To reduce protocol-dependent variation and ensure compatibility across library preparation methods (poly A vs. Total RNA), we applied a multi-step gene filtering procedure: (1) Low-expression filtering: We excluded genes with a maximum counts-per-million (CPM) value below 5 across all samples. (2) Ranking and normalization: We ranked expression values for each gene and converted them to percentiles within each protocol group. (3) Detection of protocol-sensitive genes: We excluded genes if the median percentile shifted by at least one quartile ( ≥ 25 percentile points) between protocols. To assess distributional overlap, we calculated the 5^th^ and 95^th^ percentile expression values for each gene within each protocol-defined sample class. We excluded genes if the 95^th^ percentile in one protocol was lower than the 5^th^ percentile in the other, indicating a lack of overlap. We applied this criterion in both directions (polyA vs. Total RNA). Additionally, we removed genes if 95% of expression values in one protocol consistently exceeded — or fell below — 95% of values in the other protocol.

To further ensure compatibility with earlier genome builds and improve annotation reliability, we excluded 1283 genes present only in GENCODE v31 and removed all histone genes. After applying all filters and restricting the dataset to protein-coding genes, we retained a final set of 17,061 genes for downstream analysis.

### Quantile Normalization and Batch Correction

We trained a quantile normalization model on the solid tumor (n = 456) and leukemia (n = 1313) training samples. To prepare the data, we added a value of 1 to the dataset and converted the matrix to the log2 scale. We then then applied the trained model to the entire data matrix. Next, we computed the mean across samples and saved the rank ordering of the means. We performed quantile normalization using the qnorm package in Python, which takes the raw data matrix, and the saved rank ordered means as input and returns the quantile-normalized matrix.

Using the quantile normalized matrix, we performed frozen surrogate variable analysis (fSVA) after training an SVA model on the training samples [38]. fSVA extends standard SVA by removing unwanted variation from new samples using adjustment factors learned from the training dataset. We first applied SVA to the training set to identify surrogate variables, which represent latent factors capturing batch effects. The algorithm then “froze” the relationship between these surrogate variables and the expression features by storing the projection matrix.

For each new sample, fSVA used this projection to estimate surrogate variables, regressed them out of the sample’s expression profile, and produced variation-corrected data directly comparable to the training set.

### Principal Component Analysis and Feature Selection

We assessed the discriminatory power of principal components (PCs) in differentiating tumor classes by performing principal component analysis (PCA) on the solid-tumor training set. This analysis generated multiple features sets, each capturing different proportions of variance (PCA65, PCA70, PCA75, PCA80, PCA85, PCA90, PCA95, and PCA99). For each dataset, we applied one-way analysis of variance (ANOVA) to the PC scores, grouping samples by their corresponding solid tumor type. We calculated the resulting F-statistics and associated p-values for each PC to quantify its ability to distinguish between tumor classes and adjusted the p-values for multiple testing using the Benjamini–Hochberg false discovery rate (FDR) correction.

We ranked the principal components according to their adjusted p-values and found that PCs 1–9 consistently showed the most significant discriminatory power across all feature sets. (Supplementary Table 6). Pairwise scatter plots of the top nine PCs demonstrated clear class separation, with distinct clustering patterns observed across tumor types (Supplementary Figure 1).

### Bootstrap and Out of Bag Error Estimation for Optimal Feature Selection

We performed bootstrap aggregation by creating two independent sets. The bootstrap set contained the “in-the-bag” data by sampling with replacement, and the number of samples matched the size of the training sets for ST (n = 456) and HM (n = 1313) datasets. The out-of-bag (OOB) set contained all data not selected during sampling. We generated 300 independent bootstrapped sets, each consisting of bootstrap samples and the corresponding OOB samples, from the from the quantile normalized batch corrected (QNBC) matrix. For each dataset, we performed PCA on the QNBC matrix to capture 65% (solid only), 70% (solid only), 75%, 80%, 85%, 90%, 95%, and 99% of the variance explained. Additionally, we generated a feature set of the top 1000 most variable genes from the training datasets for ST and HM separately. We trained our models using the bootstrapped samples and evaluated performance by running the corresponding OOB samples through each trained classifier. We ultimately used the PCA feature set that explained 70% of the variance for ST and 85% of the variance for HM (Supplementary Figure 2, Supplementary Table 7).

### Probability Threshold Determination from Out of Bag Testing

We computed an overall threshold to retain 95% of ST samples and 90% of HM samples from the OOB training results. When a class contained too few samples to meet these retention levels, we set the threshold cutoff to the maximum of 0.5 and the minimum threshold observed across all samples in any minor class.

### Stacking Classifier

We enhanced predictive performance, by implementing a stacking ensemble classifier with the StackingClassifier from scikit-learn [39] (version 1.1.1) in Python3 (version 3.10.4). We selected this approach because it leverages the complementary strengths of diverse classifiers, often improves generalization performance compared to individual models. In this ensemble, we combined multiple base learners (level 0 models) and used their predictions as input features to train a final estimator (level 1 model). The final estimator learned how to optimally combine the outputs of the base learners to produce the final classification. To prevent information leakage, the StackingClassifier trained base learners in parallel on the full training data and generated inputs for the final estimator using cross-validated predictions. This internal cross-validation ensured that each training prediction for the meta-model came from a fold where the base learner had not been trained on the corresponding sample.

In our implementation, we used Linear Discriminant Analysis (LDA), Support Vector Machine (SVM), Neural Network (MLP), Multi-class Logistic Regression (LGR) and Random Forest (RNF) as base learners. We selected these models to represent a diverse range of machine learning paradigms. We chose Logistic Regression as the final estimator because of its simplicity and robustness in combining probabilistic outputs. We calibrated the SVM using scikit-learn’s CalibratedClassifierCV with cross-validation, to improve the reliability of predicted probabilities. This calibration ensured that the probabilistic outputs fed into the final estimator were well-calibrated, producing more accurate and interpretable confidence scores in the final predictions.

### Rationale for Model Selection

We selected five base learners to ensure a diverse representation of machine learning paradigms, with each model contributing complementary strengths. LDA provides a simple, interpretable linear model that performs well when class distributions are approximately Gaussian. SVM offers a powerful margin-based classifier that handles high-dimensional data and captures non-linear boundaries through kernel functions. Neural Networks serve as flexible function approximators capable of capturing complex, non-linear relationship in the data. Multiclass Logistic Regression delivers a probabilistic, linear approach to classification with strong performance on linearly separable data. Random Forests provide ensemble-based, tree models that capture non-linearities and feature interactions while maintaining robustness against overfitting.

We chose this combination to leverage the distinct inductive biases of each model, enabling the stacking ensemble to capture both simple and complex decision boundaries and ultimately improve generalization performance. The stacked ensemble produced a final class prediction with a calibrated confidence score. Supplementary Table 8 lists the specific parameters used in each base classifier.

### Gene Set Enrichment Analysis

We used ranked Gene Set Enrichment Analysis (GSEA) v.4.3.3 to identify biological processes associated with the weighted loadings from PCA. We performed the analysis using with the biological process set c5.go.bp.v2025.1.Hs.symbols.gmt.

### Identify Principal Component Features that Distinguish Sample Groups

We identified PCA Features that discriminate between two tumor types by performing a Student T-test on each PCA features, comparing training samples from tumor type 1 with those from tumor type 2. We adjusted the resulting p-values for multiple hypothesis testing using the Benjamini–Hochberg procedure. We selected PCA features with a false discovery rate (FDR) q < 1 x 10^−20^. If no features met this threshold, we applied a relaxed criterion of FDR q < 1 x 10^−15^ (Supplementary Table 9).

### Fusion Transcript Detection with Fuzzion2 Pattern-Matching Algorithm

We ran Fuzzion2 v1.4.0 (https://github.com/stjude/fuzzion2) to detect gene fusions. Detecting gene fusions plays a critical role in discovering cancer drivers and clinical oncology testing, yet most existing software tools require hours to run and often miss lowly expressed fusions. Fuzzion2 applies pattern matching to identify known gene fusions in unmapped paired-read RNA-Seq data. Given a set of patterns representing fusion transcript breakpoints, it finds every read pair matching any of the patterns. It detects both exact and inexact (fuzzy) matches, with fuzzy matching tolerating variations caused by sequencing errors, SNVs, and indels. By using a novel index of frequency minimizers, Fuzzion2 processes a sample in only minutes.

### RNA-Seq Reprocessing Pipeline for Cross-Alignment Classifier Evaluation

We evaluated the robustness of classifier performance to differences in genome build, annotation version, and aligner, by selecting 25 samples with RNA-seq reads aligned using both BWA to the hg19 reference genome (GENCODE v19) and STAR to the hg38 reference genome (GENCODE v31).

For the BWA hg19 GENCODE v19 alignments, we sorted BAM files and extracted paired-end FASTQ reads, trimmed adaptor sequences, and realigned the reads with STAR (v2.0) in two-pass mode to both hg19 (GENCODE v19) and hg38 (GENCODE v31) reference builds.

We quantified gene-level expression with HTSeq-count, using the GENCODE version corresponding to the genome build used for alignment. We then processed the resulting HTSeq count matrices from each alignment strategy through the trained CanID model. For each method, we recorded accuracy, number of scored samples, number of correct predictions, and filtered counts (predictions below the classification threshold) to assess the impact of genome build, annotation version, and aligner on classification performance.

## Results

### Overview of CanID Prediction Pipeline and Data

#### RNA-seq Preprocessing and Stacked Ensemble Classification Workflow

Our classification scheme CanID uses RNAseq feature count data for protein coding genes (Methods). We used the feature count matrix from the training data as the raw input and processed it sequentially through: 1) quantile normalization, 2) batch correction using frozen surrogate variable analysis (fSVA) [38], and 3) PCA for feature reduction. We trained a stacked ensemble model for cancer subtype classification using the information-dense PCA features. (**Figure 1**A). In the prediction phase, the trained stacked ensemble model takes the read count data for each testing sample as input, transforms it using the set of frozen preprocessing models learned in the training phase, and outputs the predicted class along with a confidence score for the prediction.

**Figure 1.**
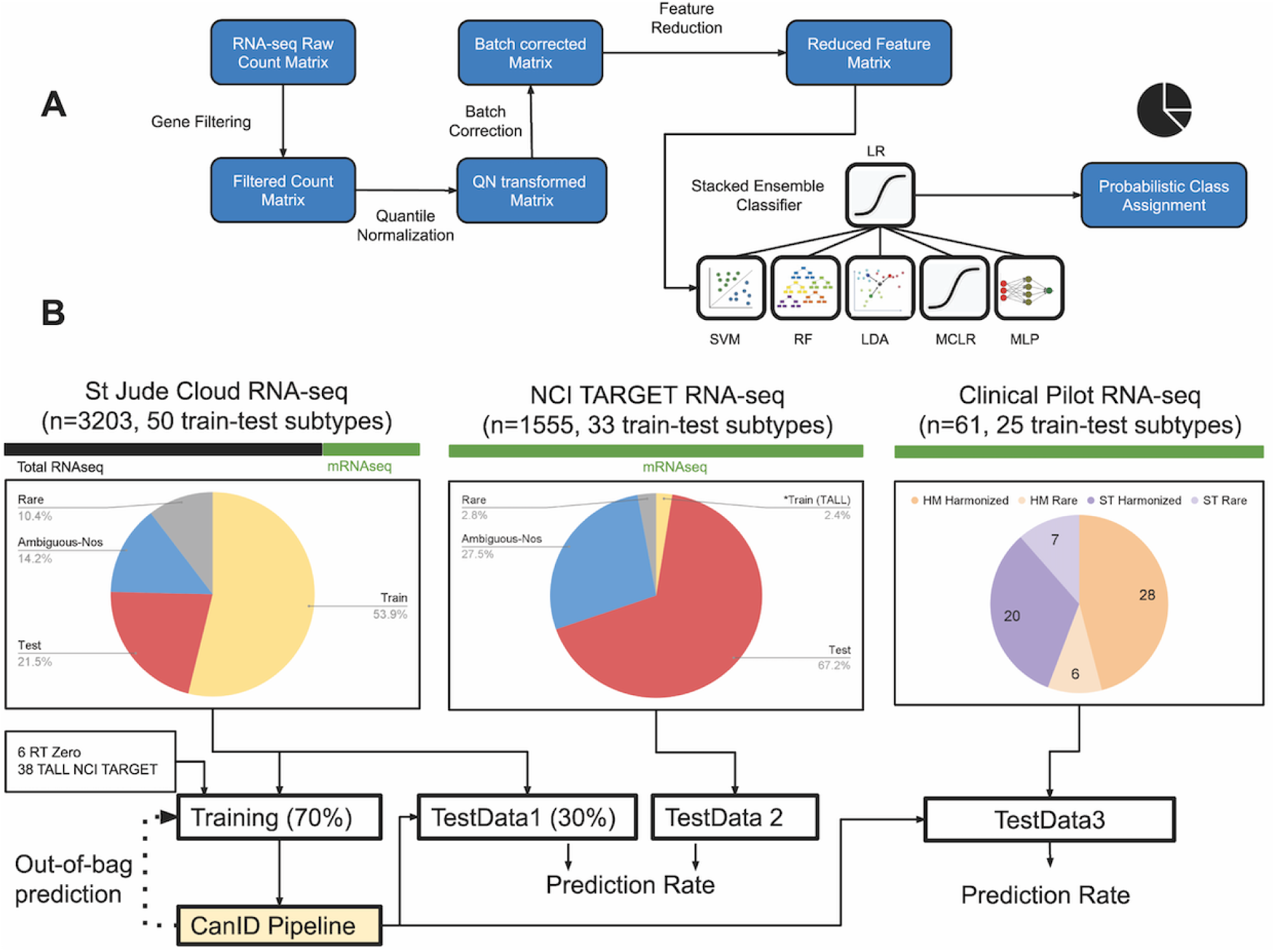
Overview of the CanID pipeline and evaluation strategy. (A) CanID pipeline: Input RNA-seq data undergo an initial gene filtering step to reduce differences between total RNA and polyA mRNA capture protocols. The filtered data are then processed through three sequential transformations: (1) quantile normalization, (2) fSVA, frozen surrogate variable analysis batch correction, and (3) PCA, principal component analysis for feature reduction. A stacked ensemble classifier is subsequently trained on the reduced PCA feature set. (B) Testing strategy: CanID was evaluated across three independent test sets. TestData1 consists of approximately 30% of SJ-Cloud patient samples (held out from a 70/30 train-test split). To ensure representation of rare subtypes, six RT, rhabdoid tumor samples were included in training, while two additional RT samples from the ZERO, zero childhood cancer precision medicine program were included in TestData1. In addition, 38 T-ALL, T-cell acute lymphoblastic leukemia samples from the NCI TARGET, National Cancer Institute’s therapeutically applicable research to generate effective treatments initiative were included in training (BCL11B = 4, NKX2 = 11, TAL2 = 6, TLX1 = 17), and 13 were held out in TestData1 (BCL11B = 1, NKX2 = 4, TAL2 = 1, TLX1 = 7). TestData2 is an independent external TARGET dataset containing 1,555 samples across 33 subtypes. TestData3 comprises an independent set of clinical pilot samples.

#### Cohorts and RNA-seq Data Sources for Model Development

As shown in Figure 1B, we used three major cohorts: 3202 RNA-seq samples (935 ST and 2267 HM) obtained from the St. Jude Cloud [23] platform, 1555 RNA-seq samples (464 ST and 1091 HM) profiled by the TARGET[11] initiative, and 61 samples (27 ST and 34 HM) obtained from a Clinical Pilot cohort that supported the development of clinical cancer genomic profiling platforms at St. Jude Children’s Research Hospital [24]. For the SJCloud cohort, we obtained subtype information for samples profiled by clinical groups from clinical diagnoses and for samples profiled by research groups from research classifications. We harmonized and curated these samples across the entire cohort, as they were collected over a ten-year period during which some diagnostic criteria may have changed (Methods). For the TARGET cohort, we derived subtype information from the TARGET data matrix or published literature. For the Clinical Pilot project, we obtained subtype information from detailed clinical records and molecular testing results.

The SJCloud cohort includes both Total RNA-seq (∼75%) and mRNA-seq (∼25%) samples profiled by 7 research initiatives and 2 clinical initiatives (Supplementary Table 5). The TARGET cohort contains exclusively mRNA-seq samples. The Clinical Pilot cohort represents a separate clinical initiative comprised exclusively of Total RNA-seq samples.

#### Assembly and Curation of Training and Validation Cohorts

We assembled the training and validation data primarily from the SJCloud cohort. To enable model development for rhabdoid tumor (RT) and several rare T-ALL subtypes, we added 8 RT samples from the ZERO Childhood Cancer project [14] and 51 T-ALL cases from the TARGET cohort: BCL11B(n = 5), NKX2(n = 15), TAL2(n = 7), TLX1(n = 24) (Tables 1–2, Supplementary Table 10).

**Table 1:** Solid tumor disease counts.

| Disease | Name | SJ-S<br>Total | SJ-S<br>Train | SJ<br>Test1 | SJ-P<br>Total | SJ-P<br>Train | SJ-P<br>Test1 | TGT-S<br>Test2 | TGT-S<br>Train | ZERO-S<br>Test1 |
| --- | --- | --- | --- | --- | --- | --- | --- | --- | --- | --- |
| ACC | Adrenocortical<br>Carcinoma | 19 | 14 | 5 | 19 | 14 | 5 | 0 | 0 | 0 |
| ARMS | Alveolar<br>Rhabdomyosarcoma | 40 | 28 | 12 | 38 | 27 | 11 | 0 | 0 | 0 |
| ERMS | Embryonal<br>Rhabdomyosarcoma | 52 | 36 | 16 | 45 | 32 | 13 | 0 | 0 | 0 |
| EWS | Ewing Sarcoma | 35 | 26 | 9 | 33 | 24 | 9 | 0 | 0 | 0 |
| HB | Hepatoblastoma | 18 | 13 | 5 | 16 | 12 | 4 | 0 | 0 | 0 |
| MEL | Melanoma | 62 | 44 | 18 | 52 | 37 | 15 | 0 | 0 | 0 |
| NBL | Neuroblastoma | 101 | 68 | 33 | 87 | 61 | 26 | 168 | 0 | 0 |
| OS | Osteosarcoma | 85 | 61 | 24 | 70 | 49 | 21 | 88 | 0 | 0 |
| RBL | Retinoblastoma | 68 | 49 | 19 | 64 | 45 | 19 | 0 | 0 | 0 |
| RT | Rhabdoid Tumor | 6 | 5 | 1 | 6 | 5 | 1 | 65 | 6 | 2 |
| SYNS | Synovial Sarcoma | 11 | 9 | 2 | 8 | 6 | 2 | 0 | 0 | 0 |
| THPA | Thyroid Papillary<br>Carcinoma | 30 | 22 | 8 | 29 | 21 | 8 | 0 | 0 | 0 |
| WT | Wilms Tumor | 102 | 75 | 27 | 76 | 54 | 22 | 130 | 0 | 0 |
|  | Total | **629** | **450** | **179** | **543** | **387** | **156** | **451** | **6** | **2** |

*Abbreviations:*
**SJ** — SJ-Cloud; **S** — sample; **P** — patient; **TGT** — TARGET.
**Test1** — testdata1; **Test2** — testdata2.

**Table 2:**
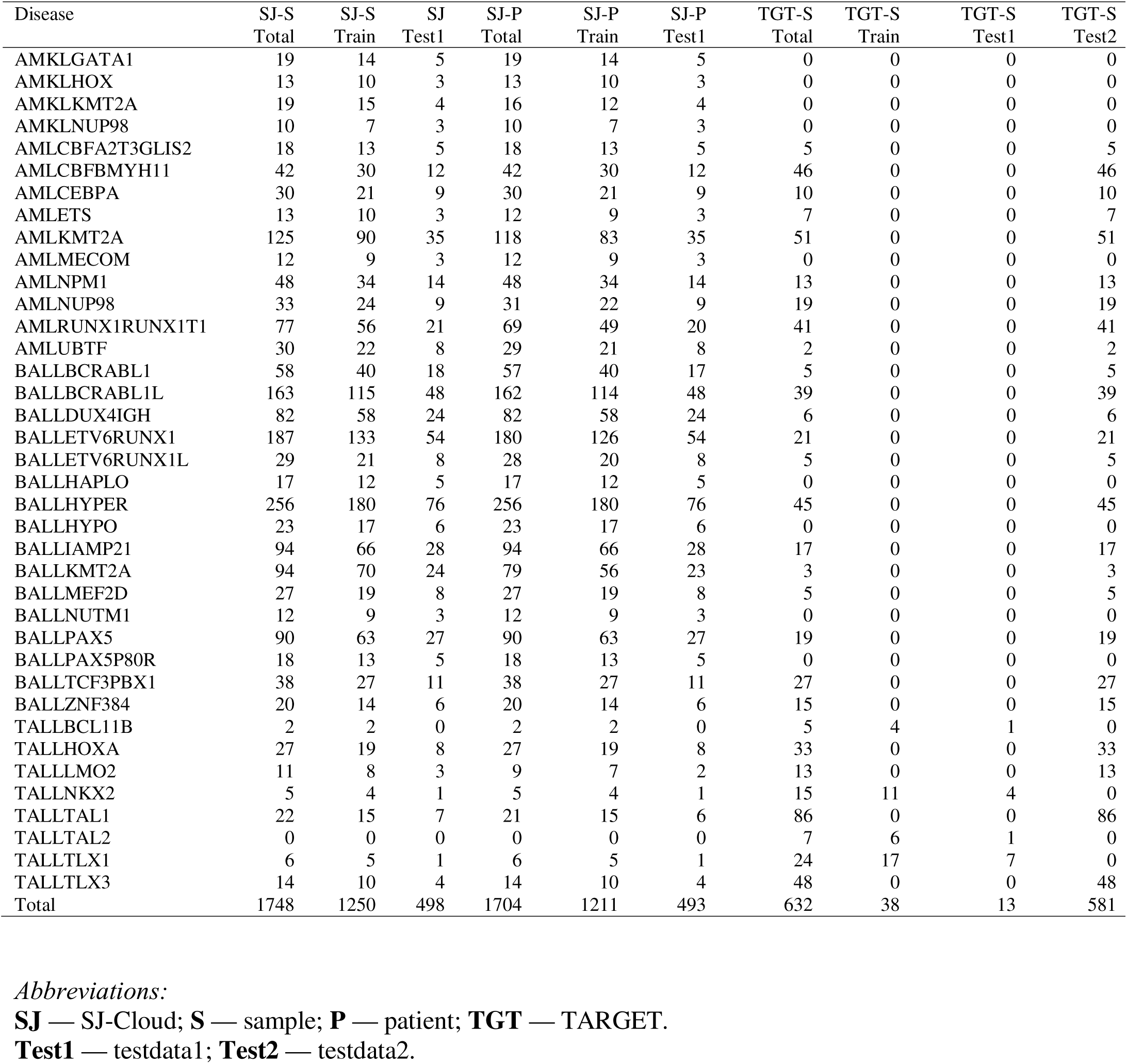
Hematologic malignancy disease counts.

We excluded the following categories of samples from model training: 1) 367 samples (16 ST and 351 HM) categorized as ambiguous or not otherwise specified (ambiguous-NOS, Supplementary Table 11) – a designation for samples that could not be definitively classified into any known disease subtypes, 2) 36 ST samples categorized as rhabdomyosarcoma (RMS), NOS – RMS samples that could not be further subclassified (Supplementary Table 12), 3) 52 ST samples that we considered low confidence after pathologist review (Supplementary Table 13) and 334 samples (202 ST and 132 HM) that belonged to rare diagnostic categories with too few samples (n < 10) to build a classifier (Supplementary Table 14). To avoid potential information leakage, we placed multiple samples from the same patient in either the training or validation set but did not split them across both groups. We used qualified samples from 70% of the patients for training which included 456 ST (13 subtypes) and 1313 HM samples [AML (14 subtypes), B-ALL (16 subtypes), T-ALL (8 subtypes)]. We reserved samples from the remaining 30% patients for validation (TestData1), including 181 ST and 522 HM samples (Tables 1–2, Supplementary Table 15).

#### Independent External and Clinical Testing Datasets

TestData2 contains the TARGET cohort using samples not included in training and serves as an independent validation cohort (Tables 1–2). We assigned 451 ST, and 581 HM samples to TestData2. We used this dataset to test the generalizability of the classifier since the TARGET data are an external data set (Supplementary Table 16). TestData3 contains the Clinical Genomics Pilot cohort with 20 harmonized ST samples (8 subtypes), and 28 harmonized HM samples (17 subtypes) We assigned one sample, SJAML030025_D1, to the rare category due to its ambiguous subtype annotation. (Supplementary Table 17). We assigned the remaining samples from the Clinical Genomics Pilot cohort to the rare group (7 ST, 6 HM, Supplementary Table 18). We included this pan-cancer clinical set to provide an independent comparison between the results of CanID and OTTER because 1) these samples were not seen in the training phase of either pipeline and 2) they have extensive clinical record and molecular testing results that allow us to derive the disease subtypes objectively.

### Model Construction and Testing

#### Optimized features for Solid Tumor and Hematologic Malignancy datasets

Given the vast difference between the ST and HM tumors, we built separate models for ST and HM datasets, respectively.

PCA is a widely used dimensionality reduction algorithm that transforms potentially correlated features into a smaller set of orthogonal principal component vectors. By reducing the data dimensionality while retaining the important information, PCA supports the construction of a robust classifier (Methods). Therefore, we applied PCA to the training data and transformed the input gene array (∼17,000 genes) into a smaller set of information-rich features that we used as input for our classification model.

To optimize the feature selection, we first tested various numbers of features based on variance explained. For example, PCA75 represents the feature set generated using PCA to explain 75% of the variance of the input data. Additionally, we generated a set of the top 1000 most variable genes from the input genes from the training datasets. We used bootstrapping to identify the optimal number of principal components to include in the PCA model (Supplementary Table 7). Briefly, for each feature set generated at a given percentage explained level, we trained 10 individual CanID models with bootstrapped training sets and repeated the process across 30 batches. We combined the out-of-bag (OOB) classification accuracy with the model complexity (number of features retained), to select the PCA feature sets that explained 80% of the variance for ST and 85% for HM, which balanced both the prediction accuracy and model complexity (Table 3; Supplementary Figure 2A). We observed that PCA95 and PCA99 showed a significant drop in performance compared to the other PCA sets used for the HM dataset, indicating that they had captured the noise in the training data (Supplementary Figure2B, 2C).

**Table 3:** Feature set accuracy in solid tumors and hematologic malignancies. For the ST, solid tumor feature sets, the principal component feature set that described 80% of the variance (PCA80) and feature set that contained the top 1000 variable genes (TOP1000) achieved the highest average accuracies. PCA80 had no significant difference between PCA PCA70, PCA75, PCA95, or TOP1000. In contrast, PCA80 significantly outperformed PCA65, PCA85, PCA90, and PCA99 at an FDR-adjusted significance threshold of 0.05. Based on this balance between model parsimony and predictive accuracy, we selected PCA80 as the optimal feature set. For the HM, hematologic malignancy feature sets, PCA85 achieved the highest average accuracy, with no significant difference compared to PCA80 or PCA90. In contrast, PCA85 significantly outperformed PCA75, PCA95, PCA99, and TOP1000 at an FDR-adjusted significance threshold of 0.05. PCA85 was therefore selected as to optimal feature set for HM.

| DATA SET | SET1 | SET2 | MEAN ACC SET1 | STDEV SET1 | MEAN ACC SET2 | STDEV SET2 | PAIRED T_STATISTIC | P VALUE | FDR ADJUSTED |
| --- | --- | --- | --- | --- | --- | --- | --- | --- | --- |
| ST | pca65 | pca80 | 0.9763 | 0.0017 | 0.9840 | 0.0021 | -7.3335 | 0.0000 | 0.0002 |
| ST | pca70 | pca80 | 0.9818 | 0.0015 | 0.9840 | 0.0021 | -2.3717 | 0.0418 | 0.0654 |
| ST | pca75 | pca80 | 0.9838 | 0.0021 | 0.9840 | 0.0021 | -0.2641 | 0.7976 | 0.8302 |
| ST | pca80 | pca85 | 0.9840 | 0.0021 | 0.9735 | 0.0032 | 8.6678 | 0.0000 | 0.0001 |
| ST | pca80 | pca90 | 0.9840 | 0.0021 | 0.9761 | 0.0046 | 6.0000 | 0.0002 | 0.0007 |
| ST | pca80 | pca95 | 0.9840 | 0.0021 | 0.9831 | 0.0046 | 0.6882 | 0.5086 | 0.5722 |
| ST | pca80 | pca99 | 0.9840 | 0.0021 | 0.9728 | 0.0067 | 5.2509 | 0.0005 | 0.0015 |
| ST | pca80 | top1000 | 0.9840 | 0.0021 | 0.9846 | 0.0021 | -0.7579 | 0.4679 | 0.5433 |
| HM | pca75 | pca85 | 0.9449 | 0.0022 | 0.9506 | 0.0021 | -5.7998 | 0.0003 | 0.0004 |
| HM | pca80 | pca85 | 0.9491 | 0.0022 | 0.9506 | 0.0021 | -1.5519 | 0.1551 | 0.1714 |
| HM | pca85 | pca90 | 0.9506 | 0.0021 | 0.9487 | 0.0040 | 2.2165 | 0.0539 | 0.0666 |
| HM | pca85 | pca95 | 0.9506 | 0.0021 | 0.7452 | 0.0080 | 89.1276 | 0.0000 | 0.0000 |
| HM | pca85 | pca99 | 0.9506 | 0.0021 | 0.5550 | 0.0074 | 154.4059 | 0.0000 | 0.0000 |
| HM | pca85 | top1000 | 0.9506 | 0.0021 | 0.9453 | 0.0030 | 4.4447 | 0.0016 | 0.0024 |

The PCA feature reduction procedure produced a relatively small set of information-enriched features. The selected PCA feature sets, PCA80 in ST (capturing 80% of the variance, 69 features) and PCA85 in HM (capturing 85% of variance, 340 features) — demonstrated strong differences among tumor types (Supplementary Figure 3) and captured biological functions essential to these tumors (Supplementary Figure 4). For example, Gene Set Enrichment Analysis (GSEA) [40, 41] applied to the gene loadings for principal component PC2 from the ST classifier revealed enrichment of immune-related signatures, including pathways associated with the adaptive immune response on the negative leading edge, and pathways related to mitotic regulation and myogenic development on the positive leading edge (Supplementary Table 19 Supplementary Figure 4). Consequently, tumors originating from developing skeletal muscle (i.e. alveolar rhabdomyosarcomas, or ARMS, and embryonal rhabdomyosarcoma, or ERMS showed higher PC2 values, whereas tumors arising from high immune cell infiltration (i.e. thyroid papillary carcinoma (THPA)), had large negative PC2 values (Supplementary Figure 3). Similarly, PC1 captured neurotransmitter transport and retina function on the positive leading edge and bone development on the negative leading edge (Supplementary Table 20; Supplementary Figure 4). Consistent with the biological processes identified, retinoblastoma (RBL) and neuroblastoma (NBL) exhibited the highest PC1 values, while osteosarcoma (OS) samples showed lower negative PC1 values (Supplementary Figure 3).

#### Classifier Architecture, Confidence Thresholding, and Performance Evaluation

CanID uses an ensemble of five individual base classifiers linear discriminant analysis (LDA), support vector machine (SVM), an artificial neural network (ANN), multi-class logistic regression (MLR), and random forests (RFS) (details in Methods). Another logistic regression model integrates the individual base-classifier outputs to produce the final prediction and confidence score. We developed a confidence threshold for the final prediction using the OOB predictions from all 300 bootstrapped models combined. CanID classified any predictions with a value below the threshold as below threshold and excluded them from valid results. We calculated the confidence score to ensure approximately 95% (ST) and 90% (HM) of the samples were above the threshold for each tumor type.

To evaluate CanID’s performance, we built a pediatric solid tumor classifier using a reference set of 456 samples from the ST dataset (450 from SJ-Cloud combined with 6 RT from Zero). We evaluated the classifier in the validation group (ST in TestData1 (N =181 with 2 RT samples from Zero) and achieved an accuracy of 0.989. CanID correctly predicted the subgroups for 173 test samples with confidence scores above the threshold (175 samples), while 6 samples fell below the threshold (**Figure 2**A, Supplementary Table 3, Supplementary Table 15).

**Figure 2.**
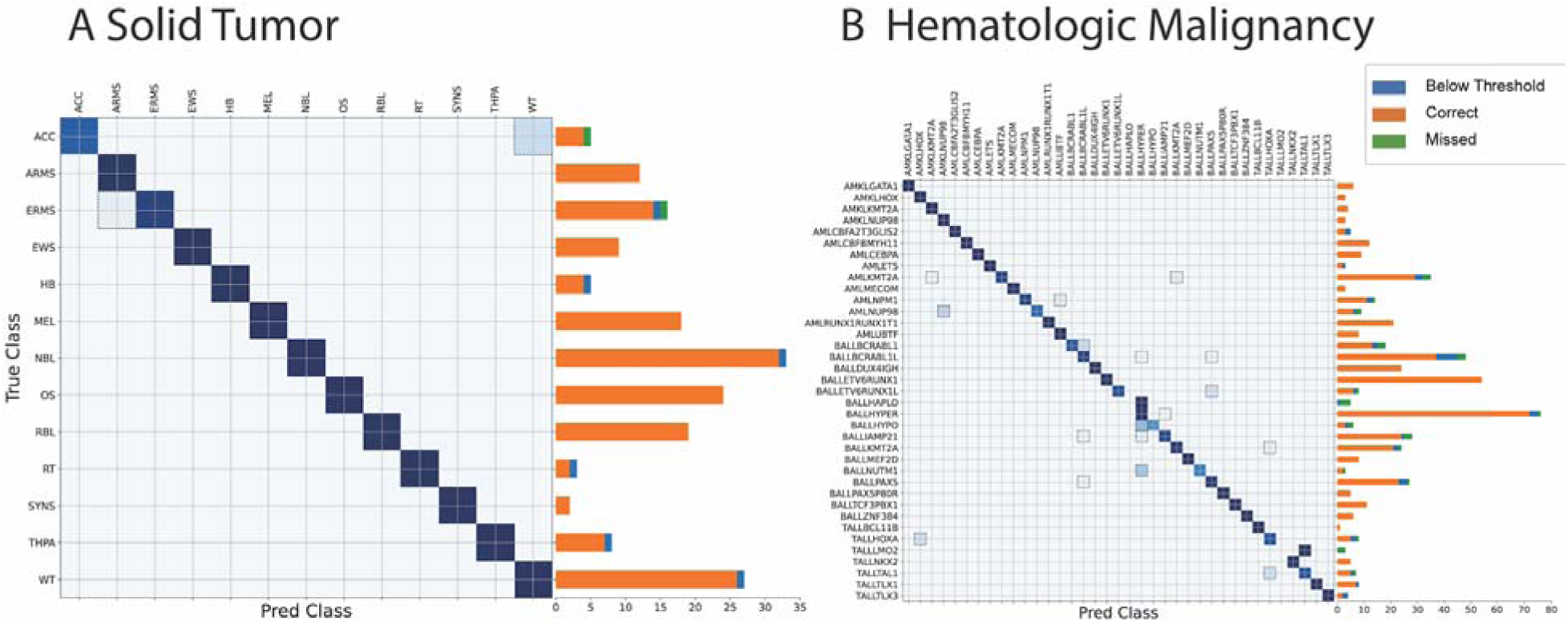
Performance of CanID on the St. Jude Cloud Test Set (TestData1). TestData1 consisted of the 30% reserved SJ-Cloud samples, supplemented with 2 RT cases from the ZERO project and 13 T-ALL, T-cell acute lymphoblastic leukemia cases from NCI TARGET (BCL11B = 1, NKX2 = 4, TAL2 = 1, TLX1 = 7). Dashed boxes highlight misclassified sample pairs. The accompanying bar chart shows the distribution of filtered samples (blue), correctly classified samples (orange), and misclassified samples (green). (A) Solid tumor dataset: overall accuracy = 0.989 (N = 181), with 6 samples below threshold. (B) Hematologic malignancy dataset: overall accuracy = 0.936 (N = 523), with 38 samples below threshold.

We evaluated the performance of the ST classifier in the NCI TARGET cohort (TestData2), an external pediatric pan-cancer dataset. CanID achieved an accuracy of 0.989 (N = 451), successfully predicting all samples above the confidence threshold (**Figure 3**A, Supplementary Table 3, Supplementary Table 16). We then evaluated performance using the ClinGen Pilot cohort (TestData3), an independent pan-cancer cohort with detailed clinical and molecular characterizations. The TestData3 set contained 20 ST samples and CanID correctly predicted the subtypes of 19 samples, while the remaining sample fell below the confidence threshold (**Figure 4**A, Supplementary Table 3). When we compared CanID with OTTER, a recently published RNA-seq based classifier for pediatric cancer [19], CanID outperformed OTTER in both sensitivity and selectivity, producing a higher number of correct predictions (19 in CanID vs. 7 in OTTER) and much lower number of incorrect predictions (0 in CanID vs. 8 in OTTER, Supplementary Table 17). Here, selectivity refers to the proportion of predictions made that were correct, reflecting the model’s ability to limit classifications to high-confidence cases.

**Figure 3.**
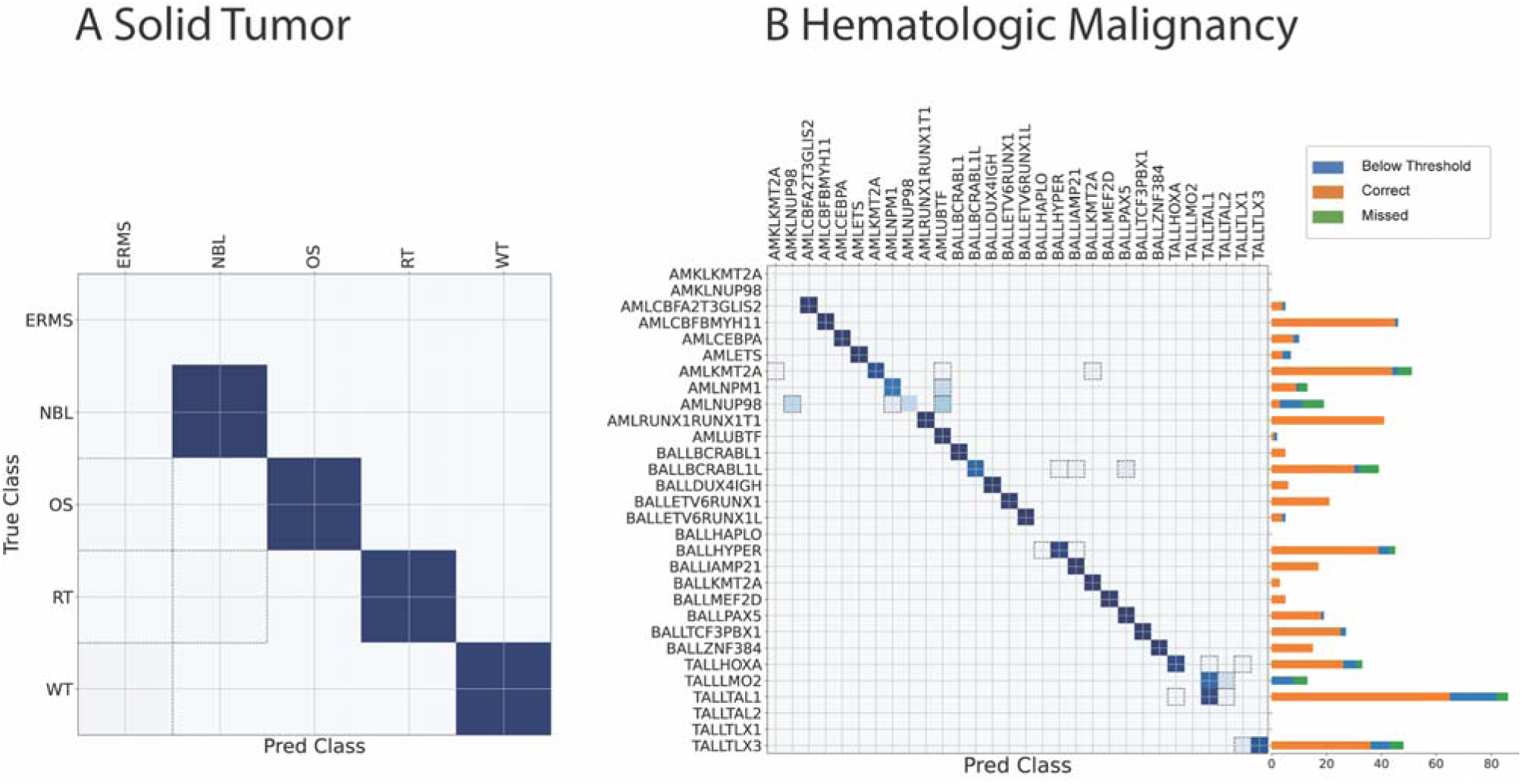
Performance of CanID on the TARGET Test Set (TestData2). TestData2 comprised all external TARGET samples. Dashed boxes highlight misclassified sample pairs. (A) Solid tumor dataset: overall accuracy = 0.989 (N = 451), with no samples below threshold. NBL achieved perfect accuracy (168/168). OS showed 98.9% accuracy (87/88 correct, 1 missed), RT showed 98.5% accuracy (64/65 correct, 1 missed), and WT showed 97.7% accuracy (127/130 correct, 3 missed). (B) Hematologic malignancy dataset: overall accuracy = 0.920 (N = 581), with 66 samples below threshold. The accompanying bar chart shows the distribution of filtered samples (blue), correctly classified samples (orange), and misclassified samples (green).

**Figure 4.**
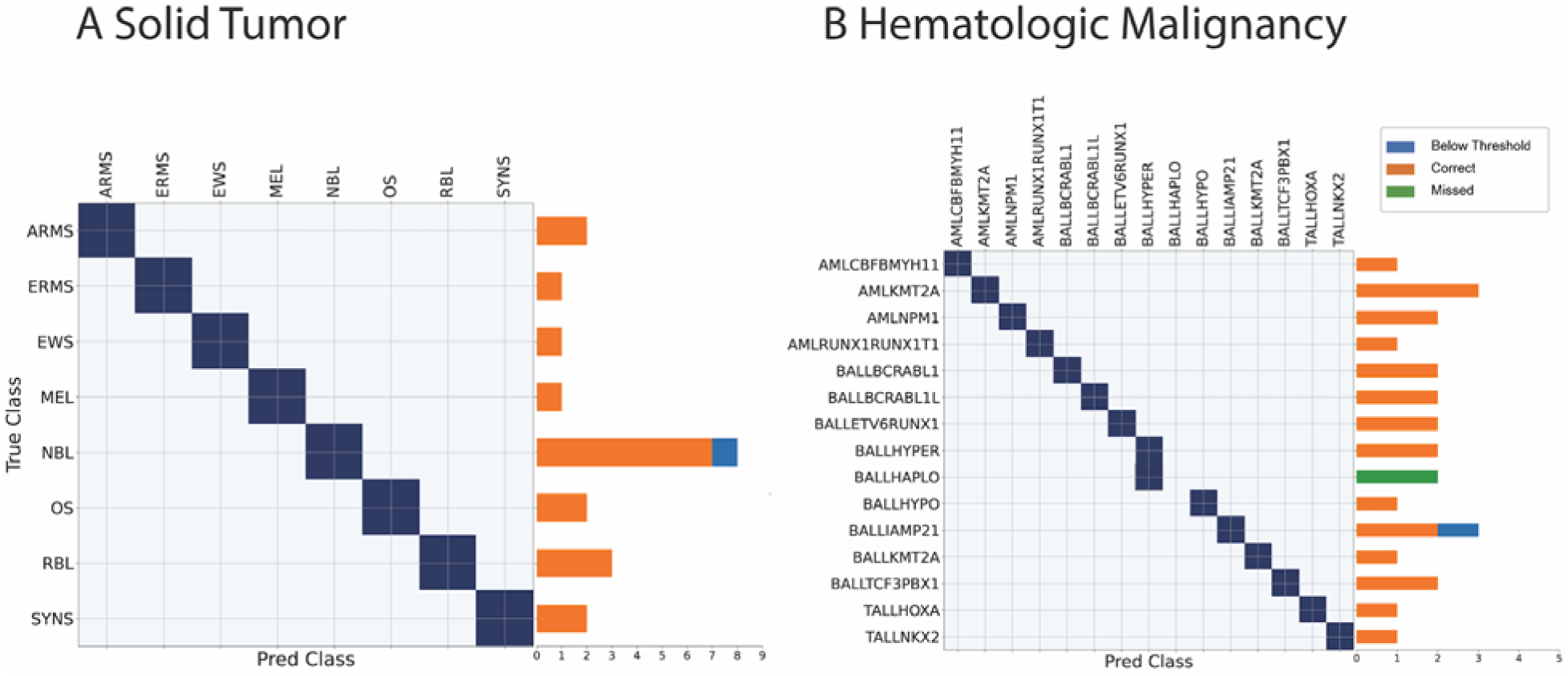
Performance of CanID on the Clinical Pilot dataset (TestData3). TestData3 consisted of Clinical Pilot samples. Dashed boxes highlight misclassified sample pairs. The accompanying bar chart shows the distribution of filtered samples (blue), correctly classified samples (orange), and misclassified samples (green). (A) Solid tumor dataset: all predictions above the scoring threshold were correct (N = 20), with 1 sample below threshold. (B) Hematologic malignancy dataset: overall accuracy = 0.920 (N = 28), with 3 samples below threshold.

We trained an HM classifier with 1313 samples. Unlike the ST dataset where tumors were derived from 13 major cancer types with largely distinct tissues, the HM dataset contained 38 related tumor subtypes from three major types. (AML with 14 subtypes, B-ALL with 16 subtypes and T-ALL with 8 subtypes). The inter-type correlation (mean 0.865 +/− 0.041) was much closer to intra-type correlation (mean 0.910 +/− 0.035) in HM compared to that of ST (inter and intra-type correlations are 0.788 +/− 0.062 and 0.876 +/− 0.061, respectively). The high expression similarity among HM subtypes increased the difficulty for classification (Supplementary Figure 5), reflected in a higher number of PCA features used in the classifier. When we tested the classifier in TestData1, it achieved an accuracy of 0.936 (N = 523 with 485 predicted and 38 below threshold, Figure 2B, Supplementary Table 15). In TestData2 (TARGET), the classifier achieved an accuracy of 0.920 (N = 581 with 515 predicted and 66 under threshold, Figure 3B, Supplementary Table 16).

Similarly, CanID achieved an accuracy of 0.920 (N = 28 with 25 predicted and 3 below threshold, Figure 4B) in the HM Clinical Pilot cohort (TestData3). It also outperformed OTTER in both the sensitivity and selectivity, producing more correct subtype predictions (23 in CanID vs. 3 in OTTER) and fewer incorrect subtype predictions (2 in CanID vs. 15 in OTTER, Supplementary Table 17).

### Subtype prediction for samples labeled as not otherwise specified (NOS)

A subset of samples from the SJCloud and TARGET cohorts (52 out of 1399 in ST and 779 out of 3359 in HM) were initially labeled as NOS. To evaluate whether CanID could classify these cases, we analyzed CanID-predicted subtype labels for approximately 70% of samples that had a confidence score higher than the subtype-specific threshold (36 in ST and 555 in HM).

*PAX5* is a critical regulator in B cell development and disrupting its gene function drives B-ALL leukemogenesis[42]. We derived a list of candidate PAX5-subtype specific marker genes by combining the top overexpressed genes (*VIPR2, ERBB2, NELL1, TPBG*) in the PAX5 subtype (vs. other B-ALLs) in the training set with those reported in literature (*PAX5, PRDM15, DENND6B, TOR4A)* [43, 44] that were also significantly overexpressed in the training set. CanID predicted B-ALL PAX5 samples in both TestData1and the NOS group and found they shared the same transcriptome signatures as those in the training set (Supplementary Figure 6).

We next evaluated the RMS, NOS subtype. The two major histological subtypes of RMS, ARMS and ERMS, have significantly different prognoses [45]. Traditionally, pathologists label RMS cases that cannot be further subtyped through histology as NOS. We analyzed 36 such samples from the SJCloud cohort and compared the predictions from CanID and OTTER. Using the ST classifier, CanID predicted 31 samples with a confidence score higher than the threshold (24 ERMS and 7 ARMS) and scored 5 samples were below the confidence threshold. OTTER generated predictions for 33 of the 36 samples: 12 classifieds as ERMS, 6 ERMS-like, 6 as ARMS, and 9 as tumor types outside the RMS category. An expert pathologist reviewed these 36 NOS samples and identified sufficient evidence to assign subtypes for an additional twelve samples (10 ERMS and 2 ARMS). Of those, the predictions made by our classifier perfectly matched the expert pathologist’s assignments, while Otter matched 6 samples, partially matched 3, misclassified 2, and made no prediction for 1 sample (Supplementary Table 12).

Our classifier’s ability to accurately subtype RMS, NOS samples is supported by signature gene expression and fusion gene analysis. *MYOG*, *TFAP2B*, *NOS1*, and *HMGA2* are well-established surrogate markers for fusion status for RMS [46]. *MYOG*, *TFAP2B*, and *NOS1* show higher expression in ARMS while *HMGA2* shows higher expression in ERMS. In both the training and testing groups, the expression pattern for *MYOG, TFAP2B* and *NOS1* matched the ARMS-specific markers previously reported in the literature. Likewise, the expression pattern for *HMGA2* in the training and testing groups matched expectations, with *HMGA2* upregulated in ERMS compared to ARMS. This pattern also appeared in the predicted subtype within the RMS, NOS group, with predicted ARMS samples showing upregulation for *MYOG*, *TFAP2B* and *NOS1*. Similarly, *HMGA2* expression was elevated in RMS, NOS samples predicted to be ERMS (Supplementary Figure 7).

Recent research suggests that recurrent *PAX3/7–FOXO1* fusions serve as the primary drivers for ARMS, and most fusion-negative ARMS are actually ERMS [47]. Therefore, we ran Fuzzion2, an in-house algorithm for detecting known fusions at high sensitivity in RNA-seq data, (details in Methods) on this RMS, NOS group. Notably, *PAX3/7–FOXO1* fusions appeared in all samples classified as ARMS (7 samples), but in none of the samples classified as ERMS (24 samples). In contrast, OTTER classified subtypes for 33 samples in this group, designating 18 as RMS and matching the fusion status (12 ERMS and 6 ARMS), producing 6 partially matched predictions (all ERMS) and 9 incorrect predictions (8 ERMS and 1 ARMS), Supplementary Table 12).

Interestingly, the *PAX3/7–FOXO1* fusion was also detected in SJRHB010463_M1, an ARMS tumor with low purity ( < 20%) [48] and one of the three unclassified samples from CanID. This finding highlights the potential for further improving the prediction performance by incorporating additional RNA-seq analyses such as fusion detection.

### Multi-omics profile of a rhabdoid tumor classified as neuroblastoma with high confidence score

TARGET sample PAKLYZ was labeled as RT, but CanID predicted it as NBL with high confidence (0.969). This incorrect prediction had the highest CanID confidence score among all other incorrectly predicted ST samples. To investigate this potential misclassification, we analyzed the whole-genome sequencing (WGS) data and expression profile of PAKLYZ in depth. Rhabdoid Tumors are driven by bi-allelic loss of the switch/sucrose non-fermentable (SWI/SNF) complex involving *SMARCB1* or *SMARCA4* [49, 50]. However, the WGS analysis of PAKLYZ from published literature [11] did not detect any mutation or copy-loss of *SMARCB1* or *SMARCA4*. This finding aligned with a previous report that assigned PAKLYZ as a *SMARCB1*-intact sample based on its elevated *SMARCB1* and *SMARC4* expression as well as clustering with normal control samples instead of RT tumors using H3K27me3 signatures [51]. Importantly, our WGS analysis identified three copy number alterations (CNAs) typically found in NBL: 1) gain of 17q, 2) loss of 1p, and 3) a *MYCN* amplicon, a well-established key driver of NBL tumorigenesis [52], with an estimated > 50 copies accompanied by *MYCN* overexpression. Therefore, evidence from the mutational landscape, the expression profile, and prior published epigenetic signature all support CanID’s classification of this case as NBL, suggesting that the initial label was likely incorrect (**Figure 5**).

**Figure 5.**
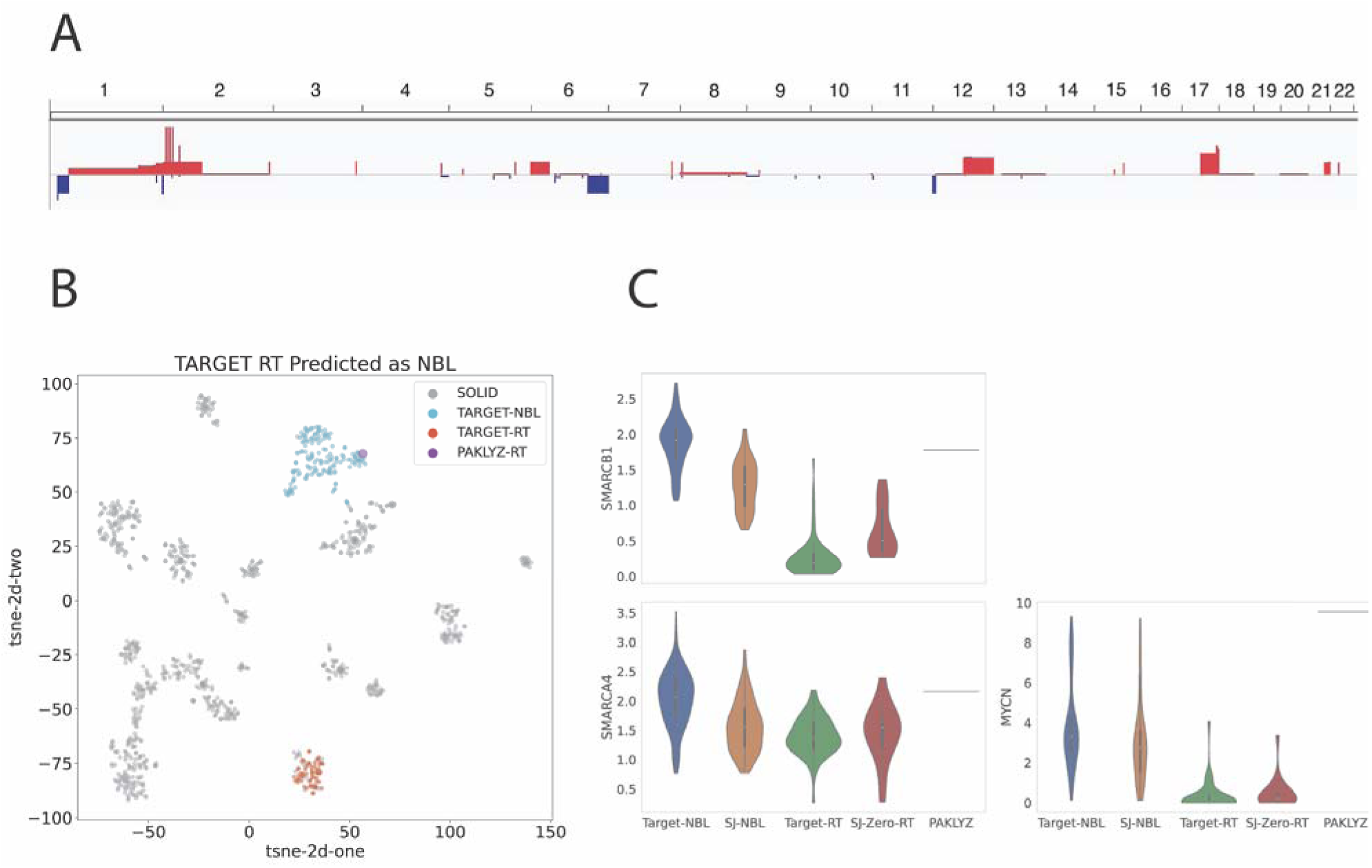
Evidence that TARGET rhabdoid tumor sample PAKLYZ is mislabeled. (A) Copy-number profile of PAKLYZ showing three signature NBL, neuroblastoma CNVs, copy-number variants: (1) gain of 17q, (2) loss of 1p, and (3) *MYCN* amplification on chromosome 2 ( > 50 copies) accompanied by *MYCN* overexpression. (B) t-SNE projection demonstrating that PAKLYZ clusters with NBL samples rather than RT samples. (C) Expression analysis of the outlier sample shows elevated *SMARCB1* and *SMARCA4* expression along with high *MYCN* expression. CanID classified this sample as NBL with high confidence (score = 0.969).

### Robustness against noise in the training samples

In the current study, the RT sample PAKLYZ from the TARGET cohort may represent an example of mislabeling in OMICs data, which can arise from human errors throughout the experimental process, including sample collection, transportation, data generation and analysis. Researchers have found that mislabeling is relatively common in published expression datasets. Using a few gender-specific markers, two studies independently identified gender-mismatched samples in 40–46% of the transcriptomic datasets, although the overall percentage of mismatched samples was relatively low [53]. In supervised learning schemes, mislabeled samples (or label noise) in the training data often deteriorate classifier performance [54]. Researchers have proposed approaches to identify and to “correct” labels for potentially mislabeled samples [55, 56]. However, without a thorough pathology review of these suspected cases, such attempts are less appropriate in tumor classification schemes. Therefore, we tested the robustness of CanID in the presence of mislabeled samples. We permutated the labels in 5, 10, 15, 20, 30, 40 and 50% of training samples and built a CanID model for each permuted training set. For each error level, we repeated the process 10 times to evaluate performance.

Although the overall accuracy of the CanID ST model decreased with the increase of label noise, the overall accuracy remained comparable to the original classifier (within 5% of the original accuracy, **Figure 6**A), with approximately 10% of the predicted sample labels falling below the scoring threshold (Supplementary Figure 8). We observed this behavior in the two large test sets with up to 30% of permuted training samples (we excluded TestSet3 due to its small sample size). The performance deteriorated substantially (lower accuracy and higher proportion below threshold) when the error level exceeded 30%. Similarly, in the more challenging HM dataset, the classification accuracy remained relatively stable with (within 5% of the original accuracy) with up to 20% label noise but declined sharply when noise exceeded 20% (Figure 6A, Supplementary Figure 8).

**Figure 6.**
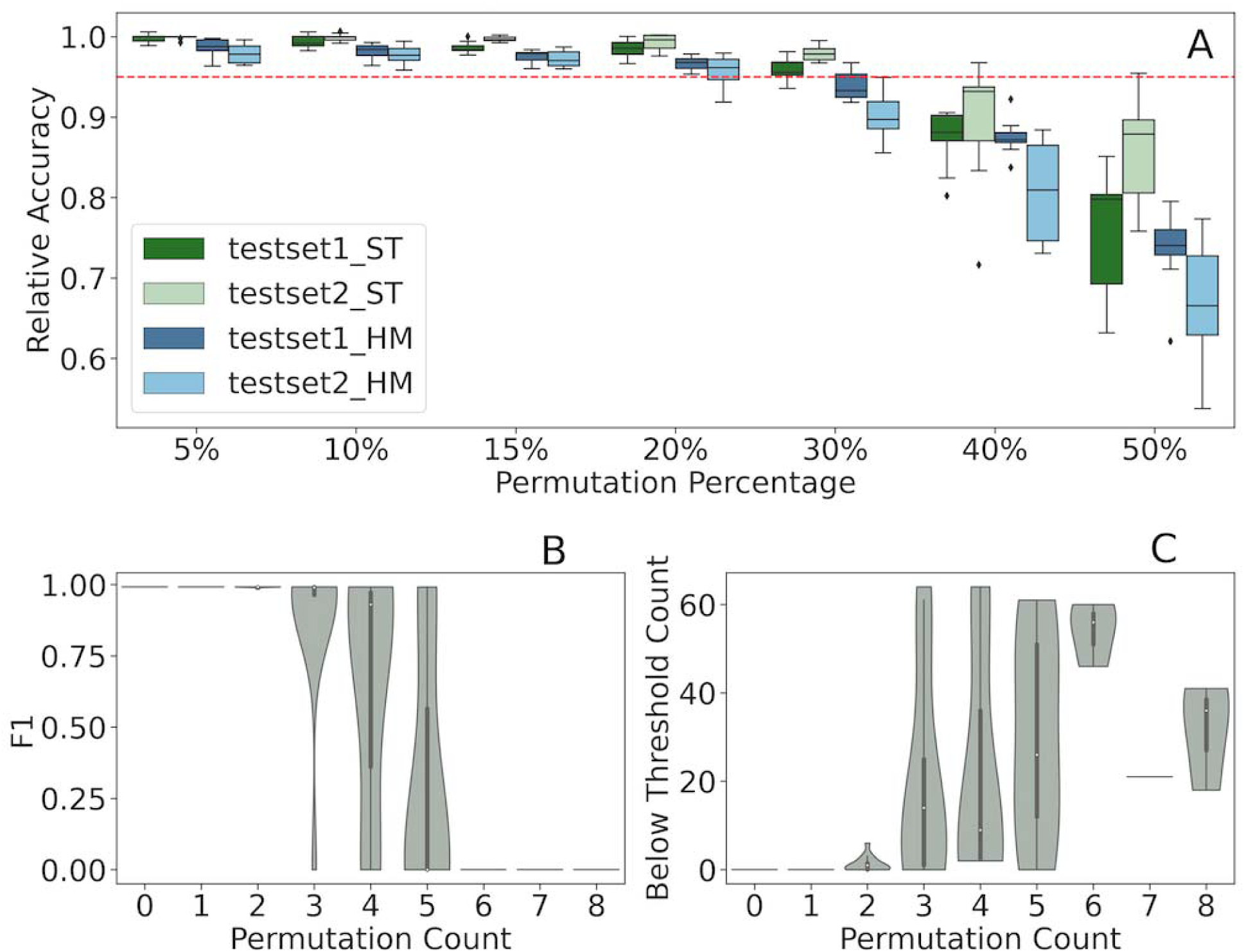
Robustness analysis of CanID. (A) Overall performance of solid tumor and hematologic malignancy CanID models on TestData1 and TestData2 under varying levels of sample label permutation, normalized to the original accuracy. The red dashed line indicates a relative accuracy of 95%. (B) Effect of RT- specific label noise on F1-score. (C) Effect of RT-specific label noise on the number of RT samples below threshold.

Real-world classification scenarios often involve both noisy labels (mislabeled training samples) and class imbalance. Mislabels in the minor classes are more detrimental because minor classes are often under-learned and the decision boundary is more susceptible to the mislabeled samples [57]. In this study, the RT class in the ST CanID model had 11 samples in the training set and 65 samples in the independently collected TARGET cohort (Testset2), providing a clear example to examine the model’s behavior in this challenging setting. We analyzed the effects on RT classification from both the overall level of noise and RT-specific noise (the number of mislabeled RT samples) for individual runs. The F1 score for the RT classification in the TARGET cohort declined strongly and significantly with increasing of RT-specific noise (β = −0.16, *p =* 5.02 X 10^−15^, linear regression) but was not affected by the overall noise level (dropped during the stepwise model selection procedure). In general, the CanID model tolerated approximately 30% RT-specific label noise without a substantial loss of performance, which is comparable to the tolerance to the overall performance (Figure 6B). Similarly, the RT-specific noise but not the overall error rate, contributed significantly and positively (β = 6.53, *p =* 1.60 x 10^−8^, linear regression) to the number of RT samples below threshold due to the confidence score dropping below the run-specific threshold (Figure 6C).

### Robustness to reference build and alignment

We evaluated the robustness of classifier performance to differences in genome build, annotation version, and aligner using human genome version 38 (hg38) and human genome version 19 (hg19). We trained CanID with reads mapped to hg38 using the STAR aligner in two-pass mode and quantified expression with HTSeq counts generated from GENCODE v31. To assess performance under alternative alignment strategies, we identified 25 samples with paired alignments – Burrows–Wheeler Aligner (BWA) to hg19 (GENCODE v19) and STAR to hg38 (GENCODE v31) – enabling within-sample comparison across reference build, annotation release, and aligner. (Methods) The dataset contained one ST case (adrenocortical carcinoma (ACC)) and multiple HM subtypes including AML (CBFA2T3-GLIS2, NUP98, GATA1, KMT2A, HOX), and B-ALL subgroups (MEF2D, PAX5P80R, ZNF384) (Supplementary Table 21). All processing methods achieved an accuracy of 1.0, with only 4–5 samples per method filtered (Supplementary Table 22). These finding indicate that classifier accuracy was unaffected by changes in genome build, annotation version, or aligner.

### Exploratory analysis of prediction for samples from rare subtypes not included in the training

A common assumption for classifiers, or supervised learning schemes is that the set of classes (or labels) encountered in deployment matches the set included in training (i.e., closed-set assumption). However, in real-world applications, most classifiers encounter classes or labels not covered in training (i.e., open-set conditions). Although specialized algorithms are preferred to address this more realistic but also challenging scenario [58–60], we evaluated the performance of our ST classifier, which has stronger inter-class separations, using samples from rare cancer subtypes.

We analyzed 202 samples in the SJCloud Rare ST cohort, covering 97 different subtypes (1–8 samples per subtype). Unlike the minimal filtering in TestData1 (3%, 6 out of 181 below threshold), CanID did not classify half of the samples from rare subtypes (52%, 105 out of 202). Among the predicted samples, we observed several patterns in which most samples from a specific subtype were predicted as a trained subtype: 1) Rare subtypes that were subclasses of an existing trained subtype (e.g. ganglioneuroblastoma predicted as neuroblastoma, botryoid-type embryonal rhabdomyosarcoma and spindle cell rhabdomyosarcoma predicted as ERMS[45], chondroblastic osteosarcoma and osteoblastic osteosarcoma predicted as osteosarcoma, etc.); 2) Rare subtypes that potentially shared a similar cell of origin with an existing trained subtype (e.g. adrenal cortical adenoma predicted as adrenal cortical carcinoma, hepatocellular carcinoma and hepatic focal nodular hyperplasia predicted as hepatoblastoma, aneurysmal bone cyst predicted as osteosarcoma, pheochromocytoma predicted as neuroblastoma, follicular thyroid cancer predicted as papillary thyroid cancer, etc.); 3) subtypes that shared similarity with an existing trained subtype (e.g., ectomesenchymoma predicted as ERMS[61], clear cell sarcoma predicted as melanoma[62], etc.).

We observed a similar pattern in the TARGET ST cohort, where 92%, (12 of 13) of clear cell sarcoma of the kidney samples fell below the threshold, with the remaining sample predicted as Wilms tumor – potentially indicating a shared cell of origin. Likewise, in the Clinical Pilot ST cohort, 86% (6 of 7) of samples were below the threshold by CanID.

Despite CanID’s overall accuracy in pediatric solid tumors, two SJ-Cloud samples from TestData1 and five TARGET solid tumor samples were misclassified with confidence scores exceeding the predetermined threshold. These included the previously discussed PAKLYZ sample (Supplementary Table 16; Supplementary Figure 9). Specifically, one ERMS sample (SJRHB031519_D1) was misclassified as ARMS; one ACC sample (SJACT030437_D1) was misclassified as Wilms Tumor (WT); three WT samples (CAAAAM, PAJMVC, PAJNVX) were misclassified as ERMS; and one OS sample (PASEFS) was misclassified as ERMS. To further understand why multiple WT samples were misclassified as ERMS, we identified PCA features with strong significant differences between ERMS and WT in the training samples – PC3, and PC4 (Supplementary Table 9, Methods). Among these differentiating principal component features, the three misclassified WT TARGET samples clustered more closely with the distribution of ERMS than with that of the WT subtype (Supplementary Figures 10 and 11). A GSEA leading-edge analysis of the gene weights on PC3 revealed strong enrichment of myogenic differentiation-related pathways — a hallmark of RMS tumors — in the negative leading edge (Supplementary Table 23; Supplementary Figure 12). Notably, the three misclassified samples were clear outliers showing negative PC3 values. In contrast, most other WT tumors exhibited strong positive PC3 values (Supplementary Figure 11, Supplementary Figure 13). Moreover, we observed elevated expression of *MYOD1* and *MYOG*, master transcription regulators in myogenic differentiation in these samples (Supplementary Figure 14). These observations suggest a possibility that these tumors represent post-therapy (chemotherapy and/or radiotherapy) tumors with induced myogenic differentiation[63].

## Discussion

We developed CanID, a general tumor (sub)type classification scheme, that uses RNA-seq feature counts as the sole input for classifying tumor subtypes. We accomplish this by establishing preprocessing models that include data normalization, batch correction, and feature reduction to transform the test data. By combining the output from five strong classifiers, we built an ensemble classification model from the transformed features to predict tumor subtypes and assign an associated confidence score to each prediction. CanID achieved high accuracy in testing samples both from the same source — split into separate training and test sets — and from independent datasets. Importantly, in a comparison with a state-of-art prediction algorithm (OTTER) using a well-characterized testing set (ClinGen Pilot), and the SJ-Cloud RMS NOS set, CanID produced better predictions for subtypes included in training and correctly identified half of the tumors from rare subtypes not included in initial training as below threshold.

Incorrect sample annotations frequently occur in transcriptomic data and pose a potential threat to classification schemes. We evaluated the performance of CanID under various levels of intentionally injected label noise (5–50%). We demonstrated that CanID models maintain robust performance with up to 30% label noise in the ST classifier and 20% in the HM classifier — exceeding the reported overall error rates observed in previous transcriptomic studies (4.1% including misclassified and unconfident samples [64], and 2.04% in cancer datasets [53]). Importantly, our analysis in RT further demonstrated that the performance for minor classes in the training phase had a similar error tolerance level.

Model robustness remains a major challenge for advanced machine learning algorithms, including deep learning. Studies have demonstrated that minor perturbations to the input of a machine learning model can cause devastating effects on its predictions [65]. In tumor classification schemes, transcriptome-based classifiers may encounter challenges from subtle differences between the two major RNA-seq platforms: the mRNA-seq protocol, which enriches mRNA with polyA tails, and the total stranded RNA-protocol, which depletes rRNAs. Despite research showing high concordance across these different RNA-seq library protocols [66], a recent study elected to exclude all samples prepared with total stranded library protocol due to apparent and consistent batch effects across tumor types [19]. However, to improve the applicability of the classifier, we aim to maintain robust performance across different library preparation protocols. By employing a feature extraction scheme that focuses on the global transcriptome pattern, CanID maintains strong performance in subtype prediction, not limited only to test cases with comparable mRNA-seq/Total RNA-seq composition as the training data. The TARGET test data generated solely by mRNA-seq, achieved comparable performance (Supplementary Table 24). Moreover, for subtypes not included in the initial training, CanID correctly labeled most of them unclassifiable. Finally, the high performance of the classification scheme in NOS rhabdomyosarcoma further demonstrates CanID’s potential to assist pathologists in determining tumor subtypes for challenging samples.

The subtype labels used for training classifiers can originate either from clinical diagnosis or from clustering analysis. While the latter approach offers the advantage of revealing potential novel subtypes that were not well characterized in current clinical practice, it also presents conceptual and technical challenges, such as confusion among distinct subtypes with similar transcriptome signatures. For example, the transcriptome profiles of near-haploid BALL samples and hyper-diploid BALL samples are highly similar, potentially due to relative overrepresentation of the same set of chromosomes [16]. Consequently, unbiased clustering may mistakenly assign them into a single cluster despite distinct clinical risk stratifications between them [67]. In the current study, we followed the World Health Organization (WHO)’s classification scheme and supplemented it with expert pathologist review of clinical and molecular testing, with the expectation that our classifiers would deliver clinically relevant performance.

In the current implementation, we explicitly use the minimal input requirement — an array of feature counts for individual genes — to maximize model portability. However, ancillary information computable from the raw sequencing files can further aid in classifying tumor subtypes in challenging samples, as demonstrated by the detection of a *PAX3–FOXO1* fusion in SJRHB010463_M1, a fusion positive ARMS sample with low tumor purity [48]. On the other side, barcode hopping remains as an in-gene fusion-based classification scheme, where multiple subtype-defining oncogenic fusions appear in a small fraction of patient samples. For example, four samples included in this study contained multiple fusions (SJBALL020141 and SJBALL020142 with MEF2D–DAZAP1 [majority] and KMT2A–MATR3 [minority], SJCBF124 and SJCBF149 with RUNX1–RUNX1T2 [majority] and CBFB–MYH11 [minority]) [68]. Since CanID relies on the whole transcriptome instead of a few “drivers”, it correctly predicted all four samples to the class defined by the majority fusion with high confidence. Therefore, both whole transcriptome-based classification schemes (such as CanID) and fusion-based classification schemes can strengthen classification power in challenging cases for the other scheme.

To integrate CanID with additional data modalities – such as mutation profiles, expression signatures, and fusion status – these inputs must be first transformed into model-ready features and combined with the PCA features derived from CanID. By concatenating these categorical and continuous variables with the PCA features, the classifier could leverage both broad transcriptomic variation and targeted molecular markers to refine subtype discrimination during training. This approach has the potential to improve classification accuracy, and the resolution of discrete subtypes.

The model’s consistent misclassification of the T-ALL LMO2 subgroup reflects the underlying biological similarity between T-ALL TAL1 and T-ALL LMO2 leukemias. Euclidean distance analysis of training sample expression profiles shows that T-ALL TAL1 samples more closely resemble T-ALL LMO2 samples than other T-ALL subgroups, indicating highly similar transcriptional programs. The t-SNE projection further supports this relationship, with T-ALL LMO2 samples clustering alongside T-ALL TAL1 cases rather than forming a distinct group. (Supplementary Figure 15). This clustering suggests that their global expression profiles are nearly indistinguishable in the reduced-dimensionality space used by the model. Biologically, the *TAL1* and *LMO2* genes encode proteins that form a multiprotein complex with other hematopoietic transcription factors to drive T-ALL leukemogenesis [69]. Studies have shown that *LMO2* is required for proper *TAL1* target binding, and the two factors cooperate to enforce a shared oncogenic program in both patient samples and mouse models [70–72]. Because these two subtypes are transcriptionally overlapping and occupy the same region in the t-SNE space, the model lacks the power to distinguish T-ALL LMO2 from T-ALL TAL1 tumor types.

A limitation of the current implementation of CanID is a requirement for at least 10 samples per subtype to be included in model construction. Consequently, rare tumor subtypes — such as clear cell sarcoma of kidney (CCSK) — while clinically relevant, cannot be integrated due to insufficient number of samples. We are actively expanding the case number for rare subtypes to broaden the classification cope of pediatric solid tumors. We plan to regularly update our CanID models as additional data become available. Furthermore, users can easily train their own model using the open-source code and instructions available on GitHub (https://github.com/chenlab-sj/CanID).

An important potential application of this work is the reclassification of legacy transcriptomic datasets. Many older studies relied solely on histopathology or limited molecular markers, resulting in incomplete or outdated subtype assignments. Applying CanID to these archived RNA-seq datasets would enable subtype prediction based on transcriptomic patterns captured through PCA-derived features. This approach could rapidly detect potential misclassifications, harmonize subtype definitions across cohorts, and enhance the accuracy of meta-analysis. In turn, it would facilitate cross-study comparisons and unlock the potential of underutilized archival datasets for precision oncology research.

## Conclusion

We developed a general classification scheme capable of generating classifiers for specific purposes. Applications across three pediatric cancer datasets with clinically relevant cancer type definitions demonstrated its robustness to sequencing library preparation protocols and batch effects, its portability, and its accuracy based on biologically interpretable features. The trained classifier distinguished most previously unclassifiable samples – 36 of 52 in ST and 555 of 779 in HM. With clear biological interpretability, and high prediction accuracy, our pan-cancer transcriptome-based classification scheme offers a valuable tool to support tumor diagnosis and enable clinically meaningful stratification of tumor types in pediatric patients with malignancies.

## Code Availability

CanID can be accessed on GitHub at https://github.com/chenlab-sj/CanID, along with documentation.

## CRediT author statement

**Daniel K. Putnam:** Data Curation, Formal Analysis, Investigation, Methodology, Resources, Software, Validation, Visualization, Writing – original draft, Writing – review & editing. **Alexander M. Gout:** Data Curation, Formal Analysis, Resources, Visualization, Writing – original draft, Writing – review & editing. **Delaram Rahbarinia:** Data Curation, Investigation, Resources, Writing – review & editing. **Meiling Jin:** Formal Analysis, Investigation, Resources, Writing – review & editing. **David Finkelstein:** Formal Analysis, Resources, Writing – review & editing. **Xiaotu Ma:** Formal Analysis, Resources, Writing – review & editing. **Jinghui Zhang:** Conceptualization, Formal Analysis, Project Administration, Supervision, Writing – original Draft, Writing – review & editing. **David A. Wheeler:** Conceptualization, Formal Analysis, Project Administration, Supervision, Writing – original draft, Writing – review & editing. **Larissa V. Furtado:** Conceptualization, Formal Analysis, Project Administration, Writing – original draft, Writing – review & editing. **Xiang Chen** Conceptualization, Data Curation, Formal Analysis, Funding Acquisition, Methodology, Project Administration, Supervision, Writing – original draft, Writing – review & editing.

## Acknowledgements

This study was supported, in part, by National Cancer Institute (NCI) grants R01CA262790 and R01CA266600 to X.C. The study is also supported by American Lebanese Syrian Associated Charities (ALSAC).

We would like to thank Andrew Thrasher for help with explaining the gene filtering process, Steve Rice for analyzing gene fusions on the rhabdomyosarcoma NOS cases, and Sarah August for manuscript editing.

## Supplementary Material

### Supplementary Figures

**Supplementary Figure 1.**
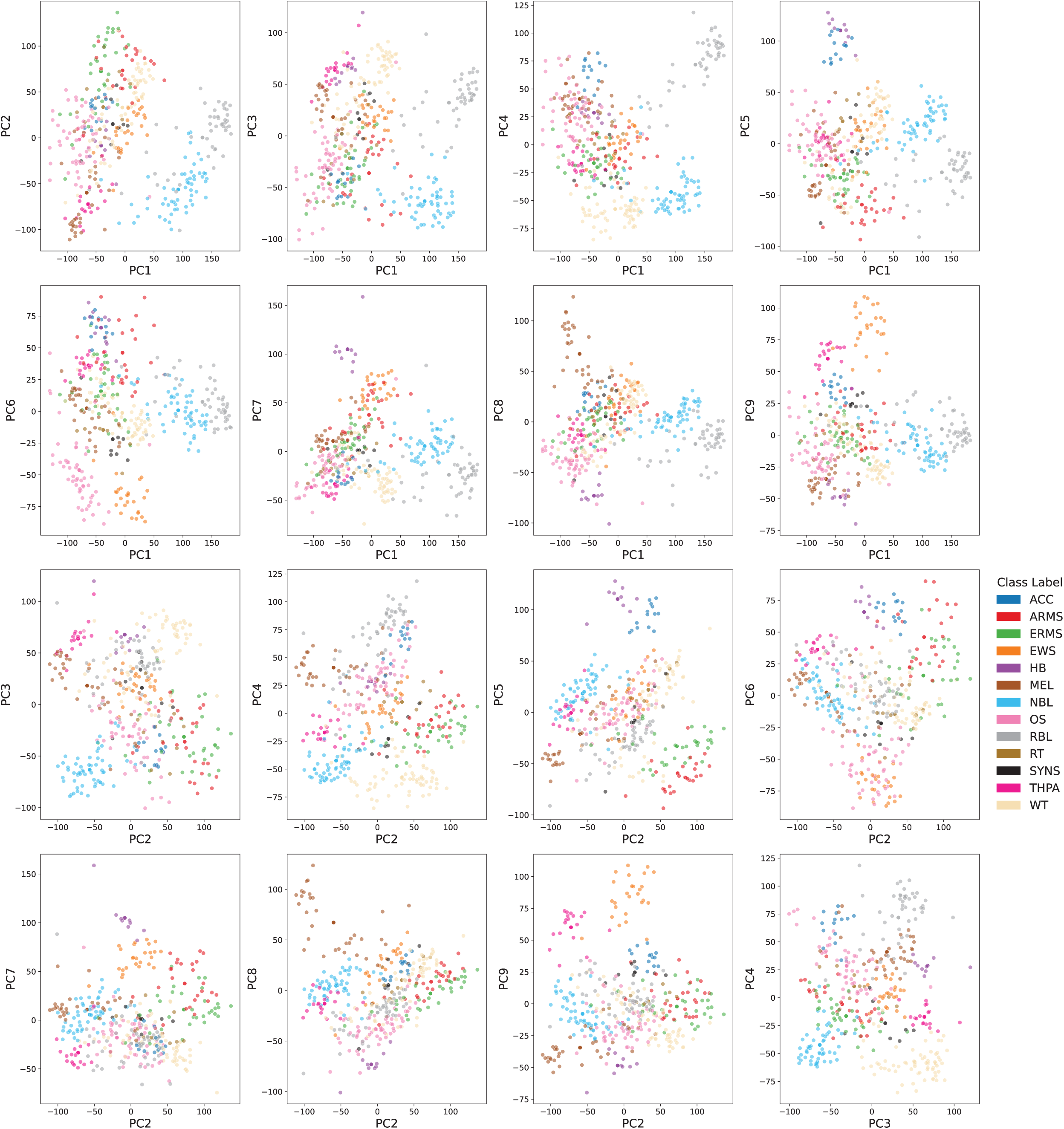
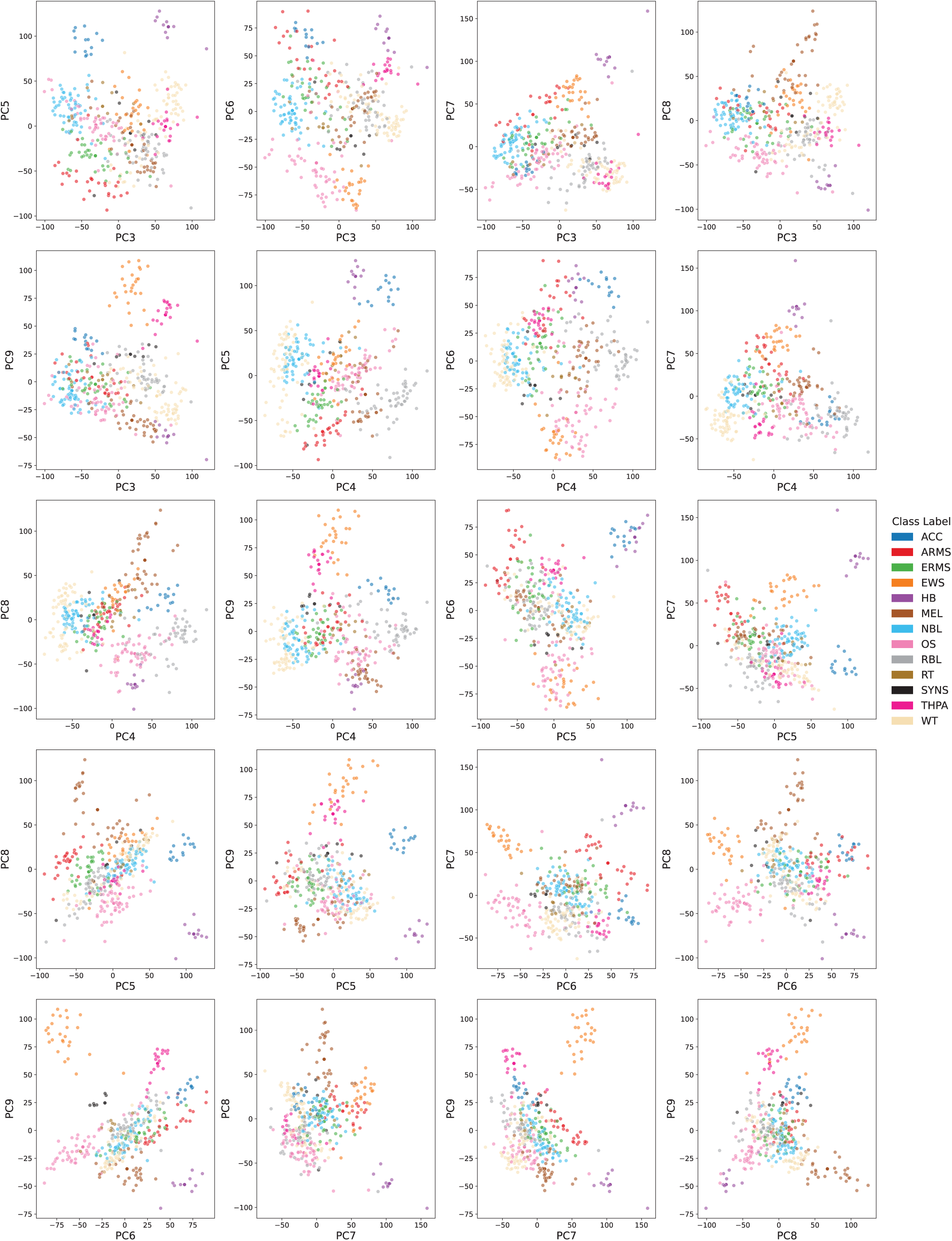
Pairwise scatter plots of the top nine principal components (PCs) illustrating tumor class separation. Each point represents an individual tumor training sample, colored according to its class. The figure displays all 36 pairwise combinations of the top nine PCs. These leading PCs, identified through ANOVA-based ranking, exhibit discriminatory power, with distinct clustering patterns observed for tumor classes across multiple component pairings.

**Supplementary Figure 2.**
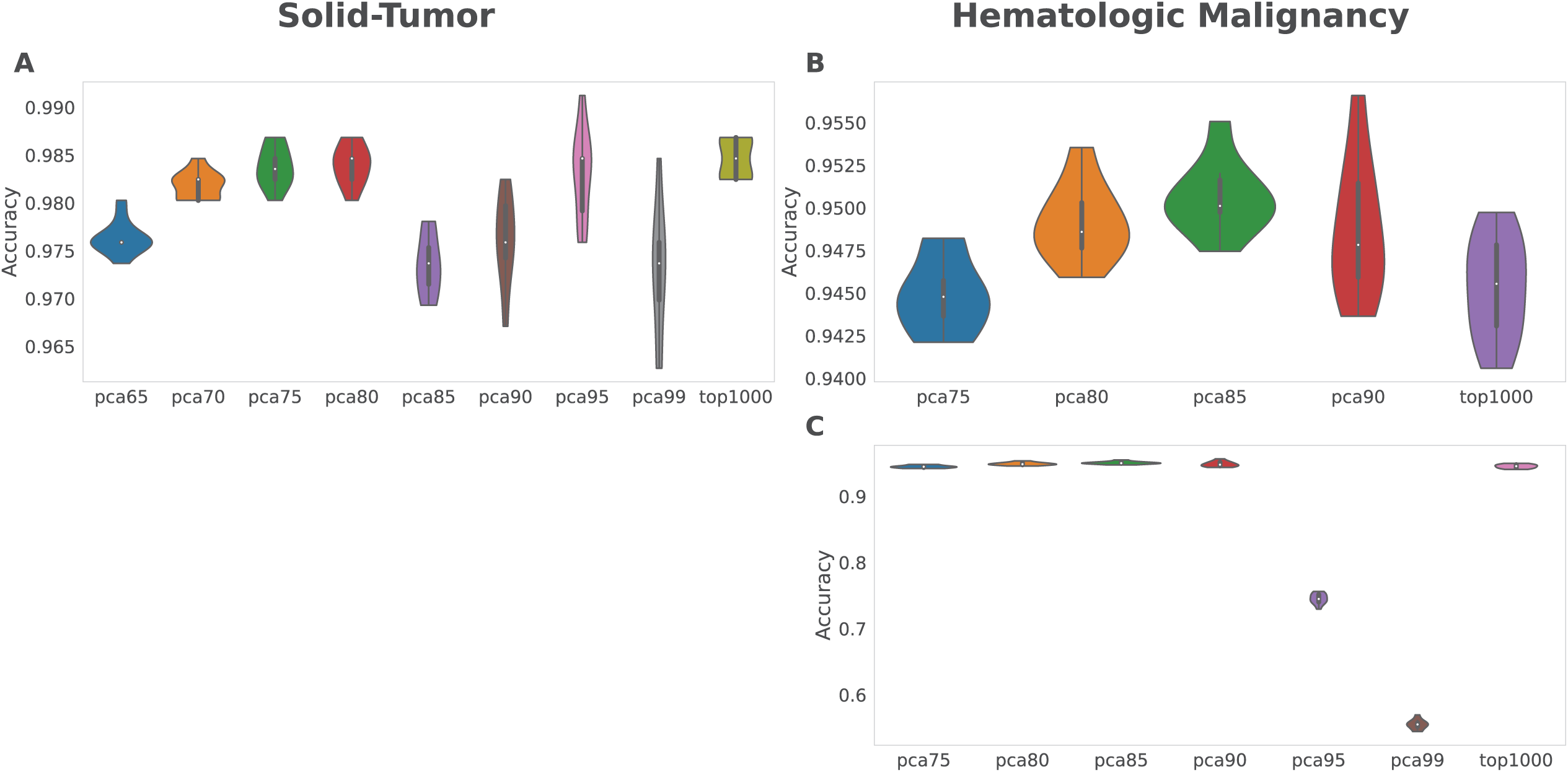
Feature set selection for tumor classification. (A) PCA80 was selected as the optimal feature set for ST, non-CNS solid tumors, while (B) PCA85 was chosen for HM, hematologic malignancies. In contrast, higher-dimensional feature sets—PCA95 and PCA99—showed a pronounced decrease in performance for HM, with mean accuracies of 0.745 ± 0.008 and 0.555 ± 0.007, respectively. (C) Comparative results across all HM PCA feature sets are shown.

**Supplementary Figure 3.**
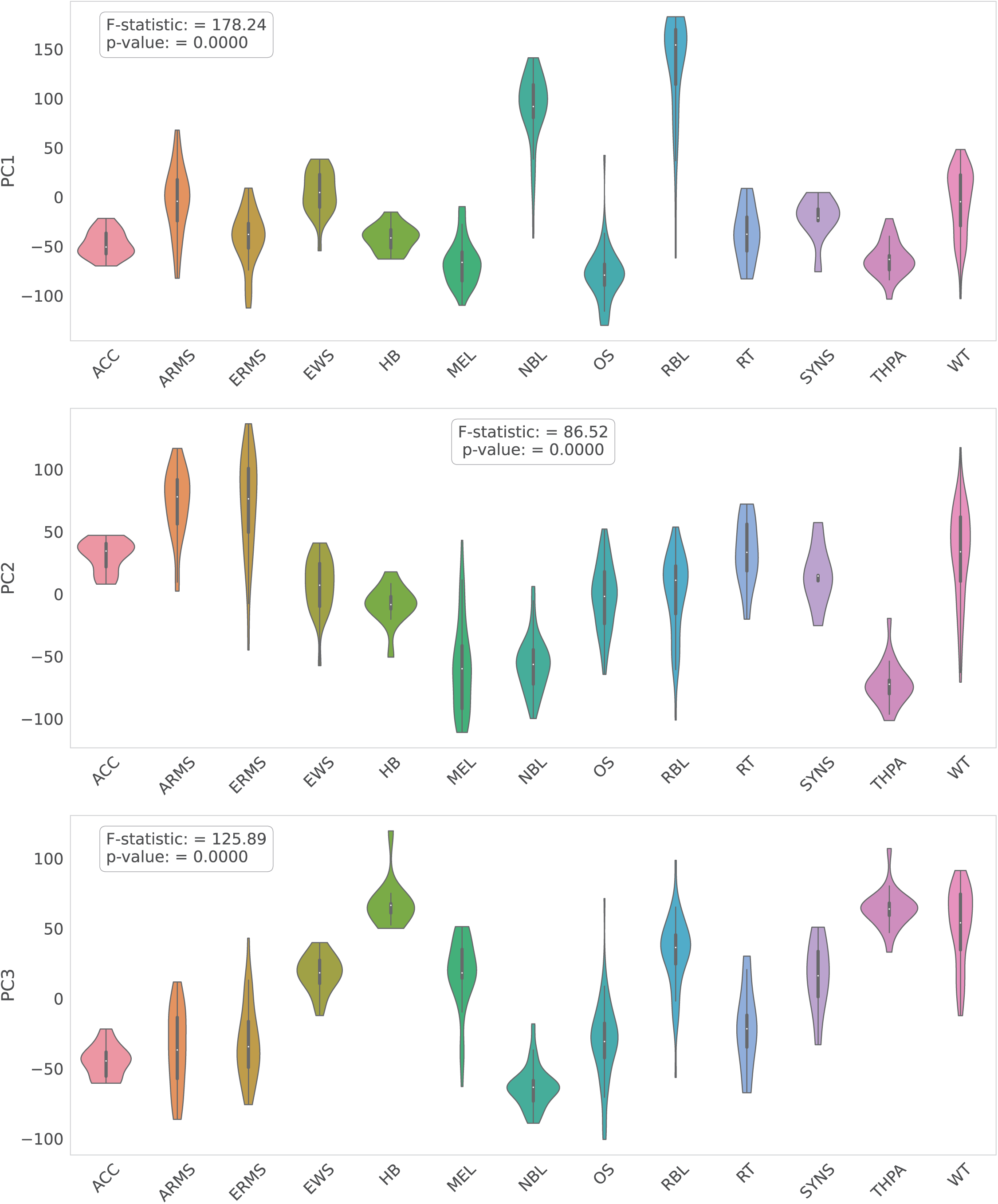
Principal Component Patterns in the Solid Tumor Cohort. Distinct tumor types are characterized by unique distributions along specific principal components. NBL and RBL exhibit strong positive PC1 values. MEL shows strong negative PC2 values, whereas ERMS, ARMS, and WT show strong positive PC2 values. Along PC3, ERMS and ARMS display strong negative values, while HB, THPA, and WT show strong positive values.

**Supplementary Figure 4.**
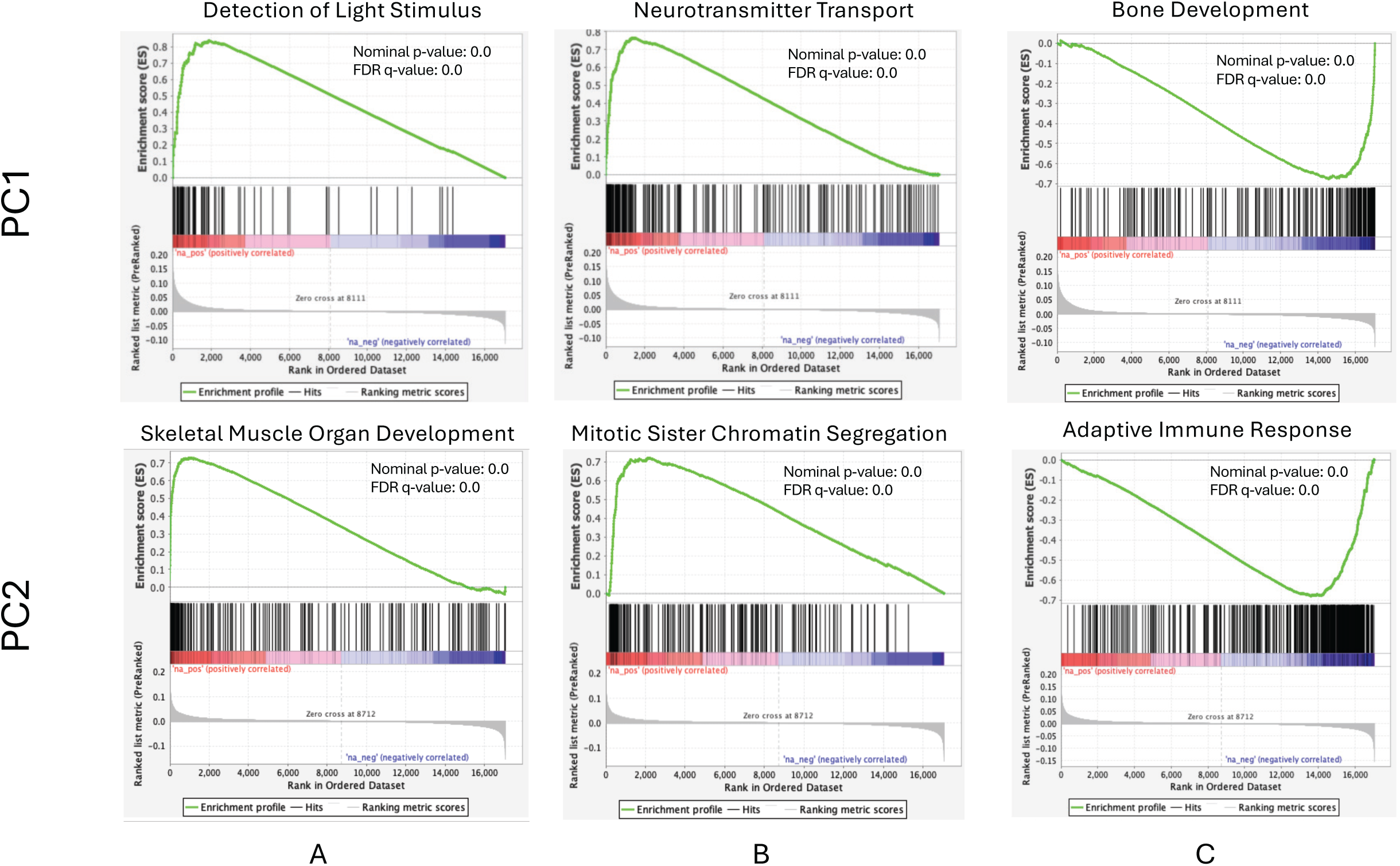
PC1 Developmental and PC2 Immune/Myogenic Signatures. Using the weighted loadings for Solid-Tumor PC1 and PC2 we performed GSEA to identify biological processes associated with genes driving these components. PC1 was enriched for developmental pathways, reflecting variation in tumor lineage and differentiation. PC2 showed immune-related signatures on the negative leading edge and mitotic and myogenic pathways on the positive leading edge. Panels A and B display enriched gene sets for up-weighted genes, while Panel C shows an enriched set for down-weighted genes.

**Supplementary Figure 5.**
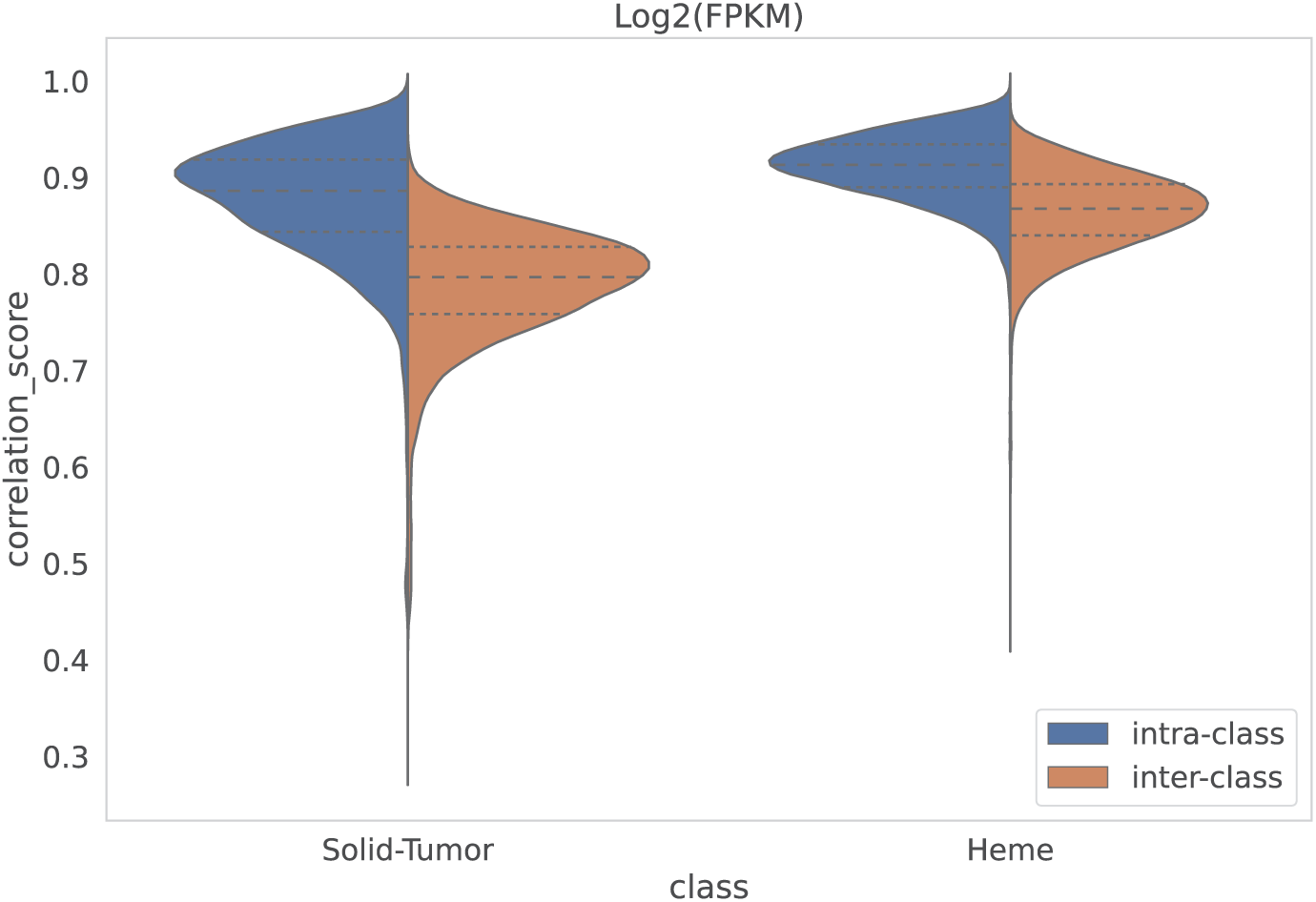
Hematologic dataset shows a narrower range of inter- and intra-class correlation scores. The solid tumor cohort exhibits an inter-type correlation of 0.788 ± 0.062 and an intra-type correlation of 0.876 ± 0.061, whereas the hematologic malignancy cohort demonstrates higher correlations, with an inter-type correlation of 0.865 ± 0.041 and an intra-type correlation of 0.910 ± 0.035.

**Supplementary Figure 6.**
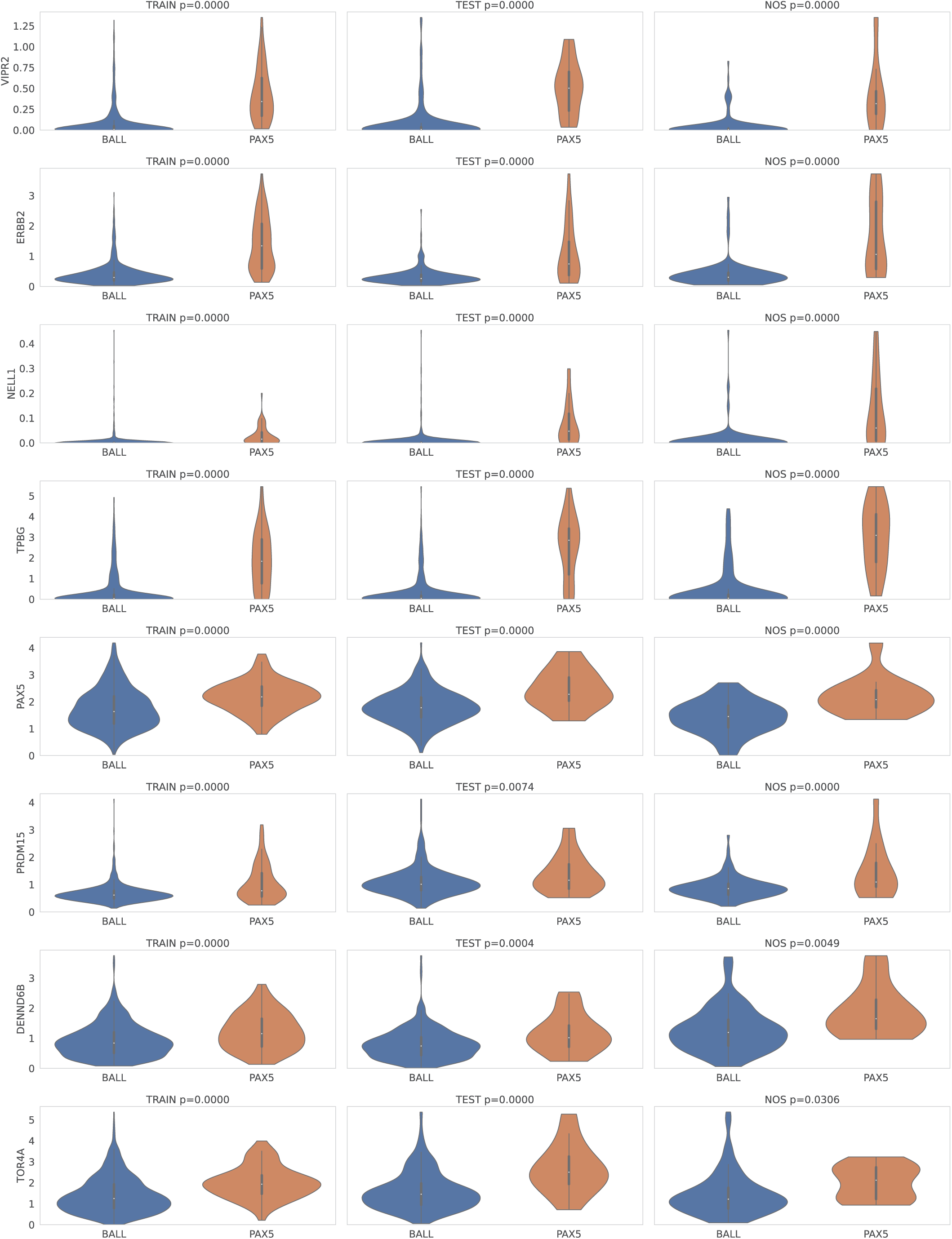
Expression Patterns of Candidate PAX5 subtype marker genes. PAX5, a key regulator of B cell development, is frequently disrupted in B-ALL, B cell acute lymphoblastic leukemia. Candidate PAX5-subtype marker genes were identified by combining the top overexpressed genes in the PAX5 subtype versus other B-ALLs in the training set (*VIPR2*, *ERBB2*, *NELL1*, *TPBG*) with literature-reported genes (*PAX5*, *PRDM15*, *DENND6B*, *TORA*). CanID-predicted subtypes (PAX5 vs B-ALL) in the NOS group mirrored the expression patterns observed in both the training and TestData1 sets.

**Supplementary Figure 7.**
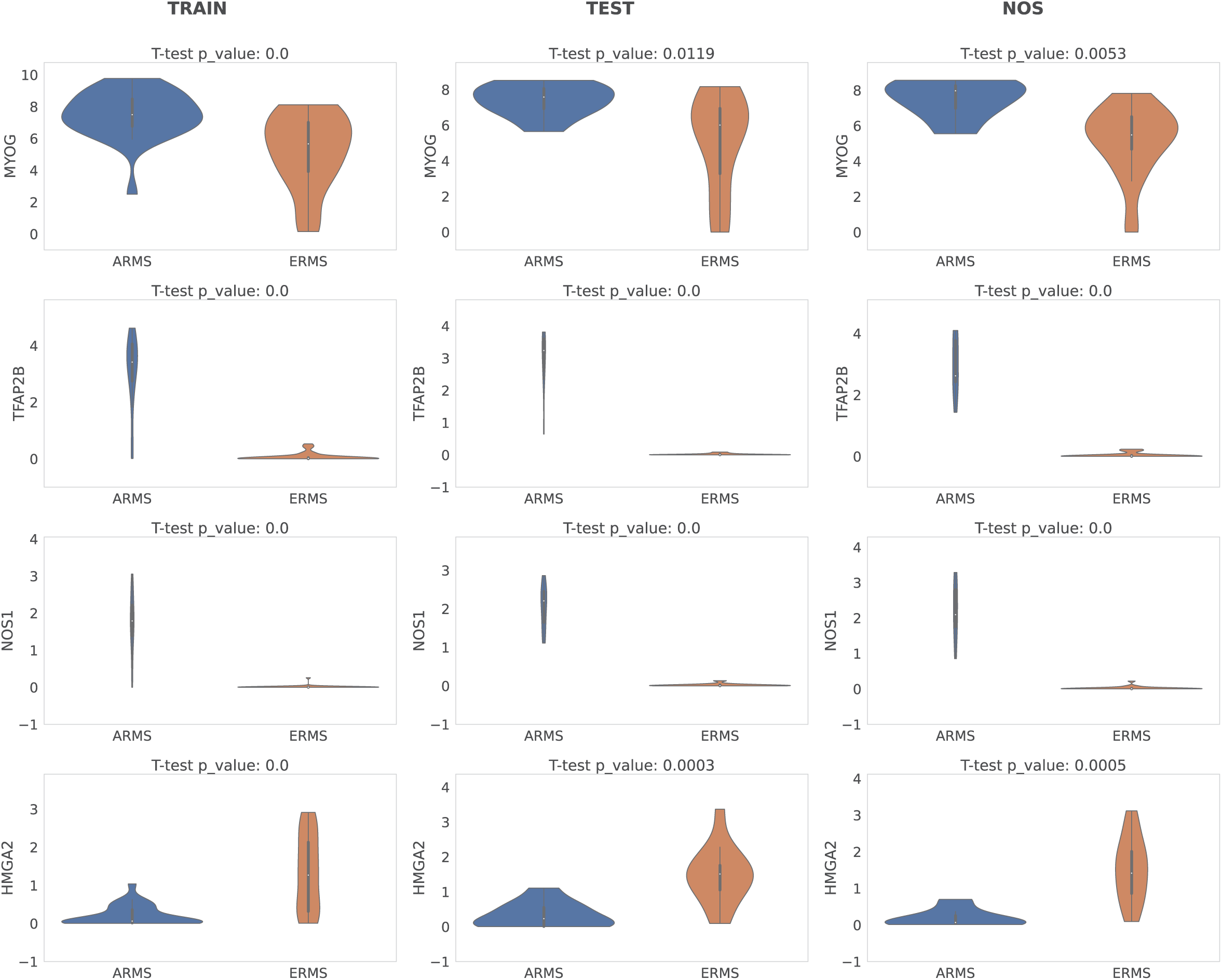
Expression pattern of rhabdomyosarcoma surrogate markers for fusion status. Violin plots of train, test, and NOS, not otherwise specified gene expression for *MYOG*, *TFAP2B*, *NOS1*, and *HMGA2*. *MYOG*, *TFAP2B*, and *NOS1* have higher expression in ARMS, alveolar rhabdomyosarcoma, while *HMGA2* has higher expression in ERMS, embryonal rhabdomyosarcoma. The CanID predicted subtype (ARMS vs ERMS) in the NOS column mirror the expression patterns seen in the train and TestData1 set.

**Supplementary Figure 8.**
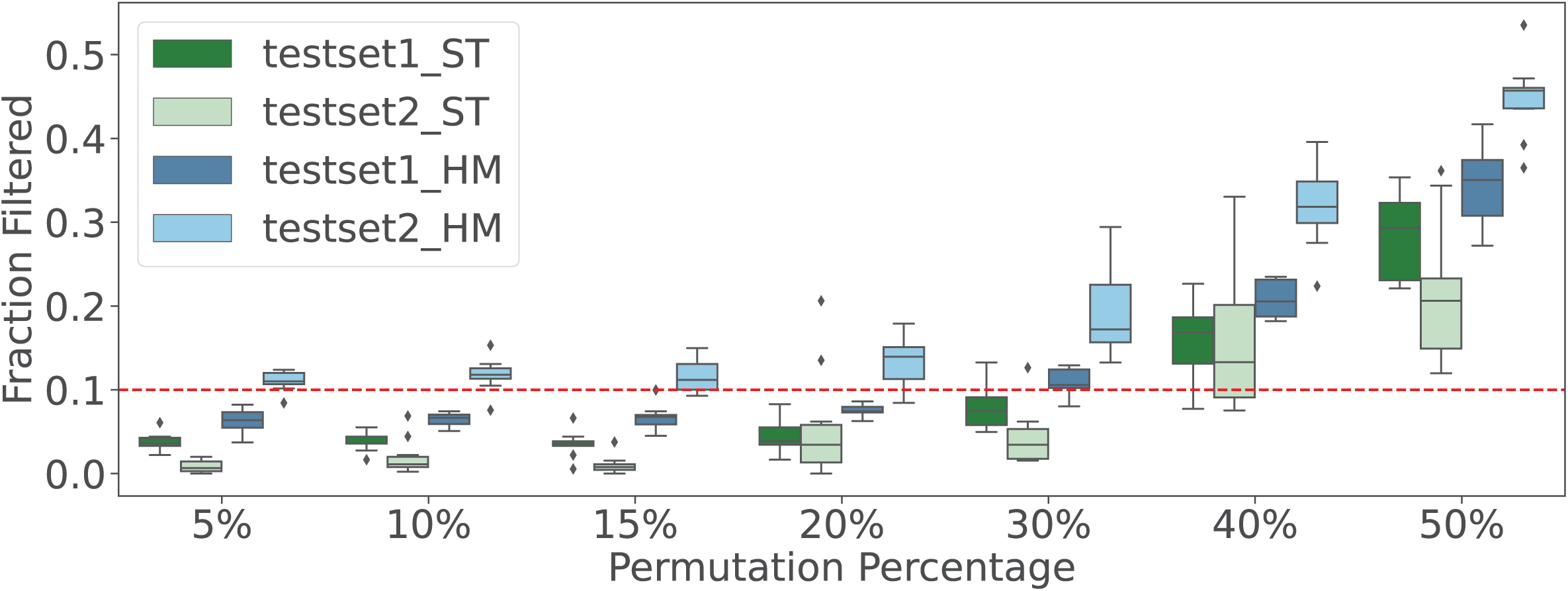
Performance robustness of solid tumor and hematologic malignancy CanID models under increasing sample label permutations. Overall performance on testset1 and testset2 is shown as the fraction of samples filtered across varying permutation percentages. The fraction filtered increases with higher permutation thresholds. The red dashed line marks a 10% filtering rate.

**Supplementary Figure 9.**
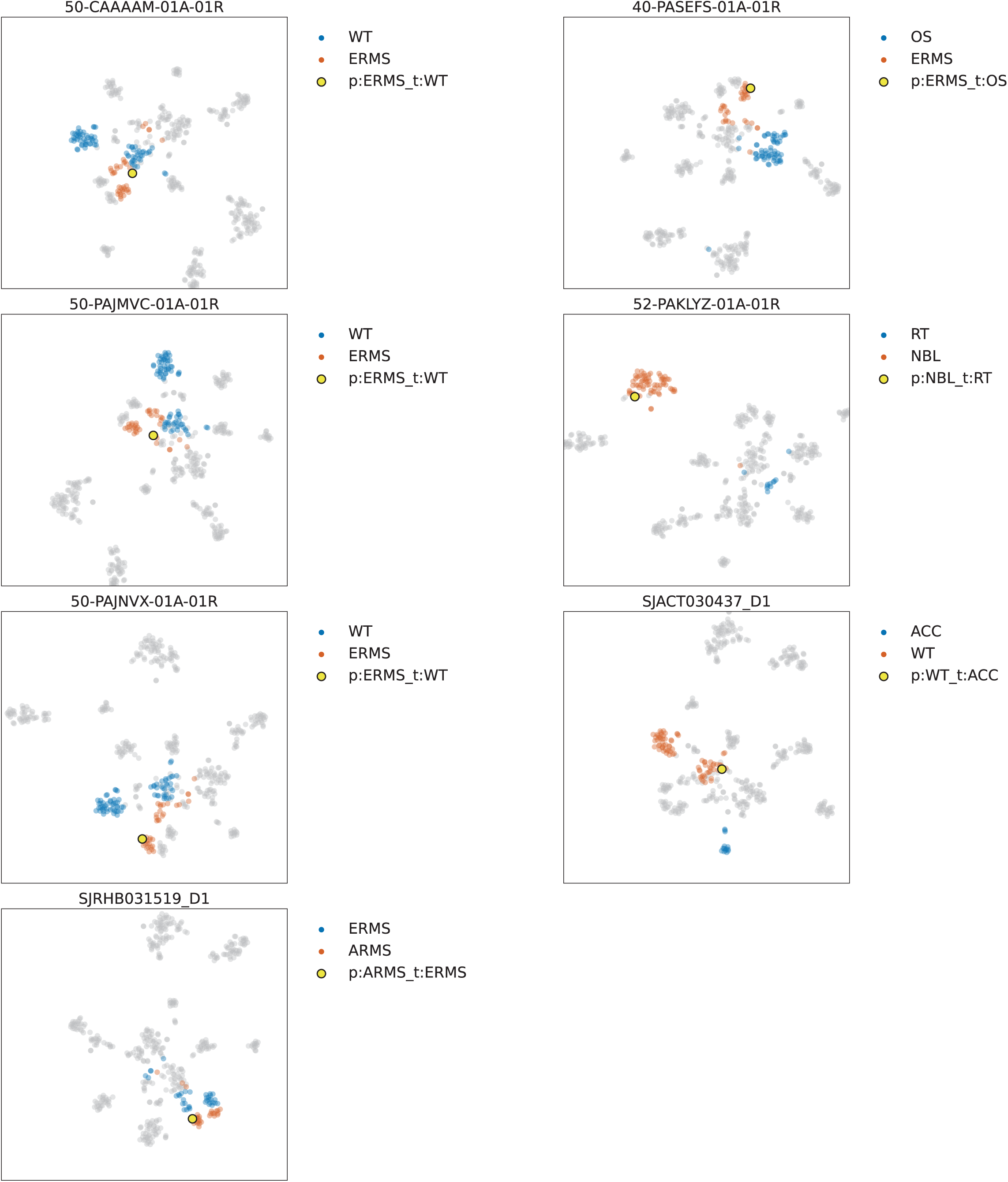
Plots of misclassified solid tumor samples cluster with their predicted type. Testset1: One ACC, adrenocortical carcinoma sample (SJACT030437_D1) was misclassified as WT, Wilms tumor and one ERMS, embryonal rhabdomyosarcoma sample (SJRHB031519_D1) was misclassified as ARMS, alveolar rhabdomyosarcoma. Testset2: Three WT samples (CAAAAM, PAJVMC, PAJNVX) were classified as ERMS, one OS, osteosarcoma sample (PASEFS), was classified as ERMS and one RT, rhabdoid tumor sample (PAKLYZ) was classified as NBL, neuroblastoma. The t-SNE projections were generated from raw counts converted to FPKM for the input gene set (N=17061).

**Supplementary Figure 10.**
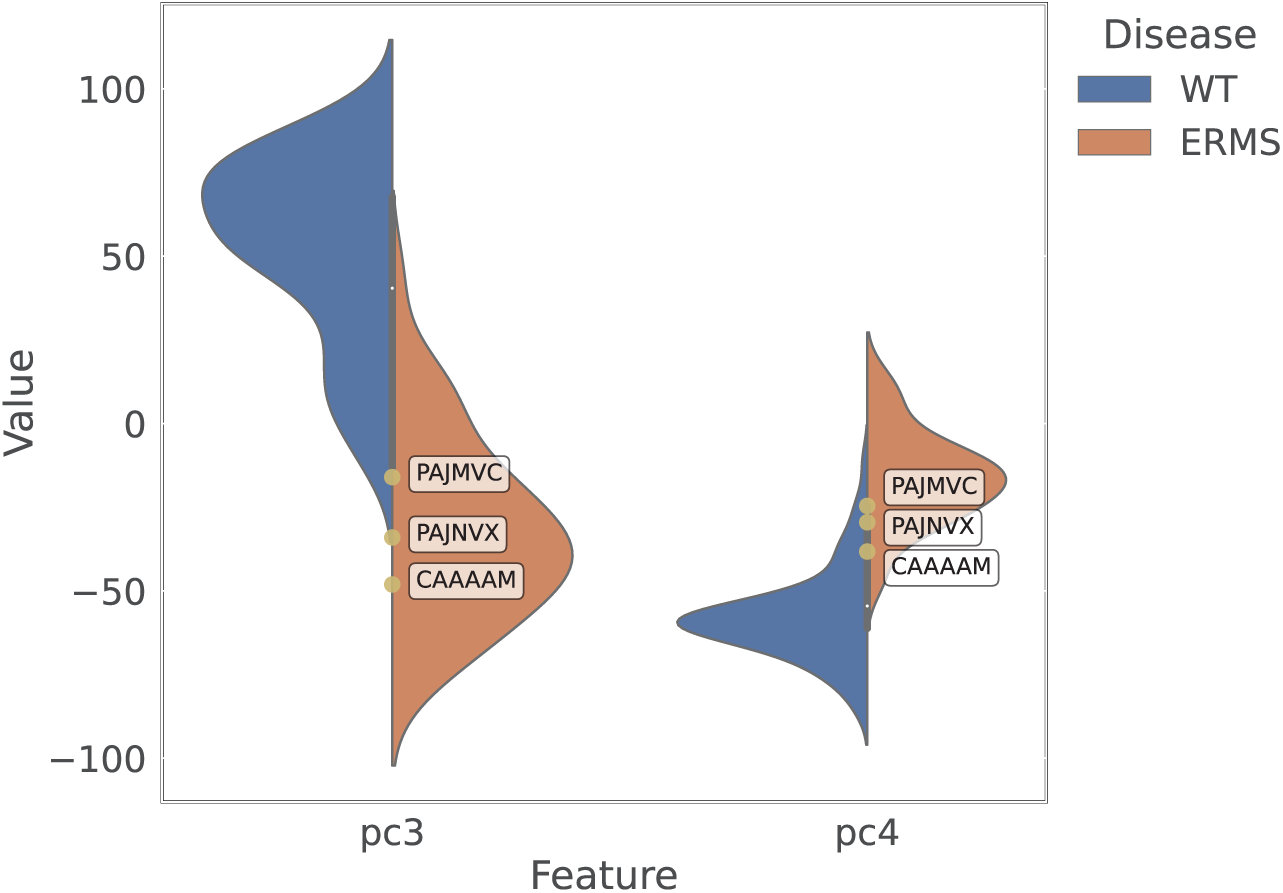
Misclassified Wilms tumor cases align with embryonal rhabdomyosarcoma profiles. Violin plots show the distributions of the principal components that distinguish ERMS, embryonal rhabdomyosarcoma (orange) from WT, Wilms tumor (blue) in the training data. Three TARGET, therapeutically applicable research to generate effective treatments samples— CAAAAM, PAJMVC, and PAJNVX—were labeled WT but predicted as ERMS and are shifted toward the ERMS distribution.

**Supplementary Figure 11.**
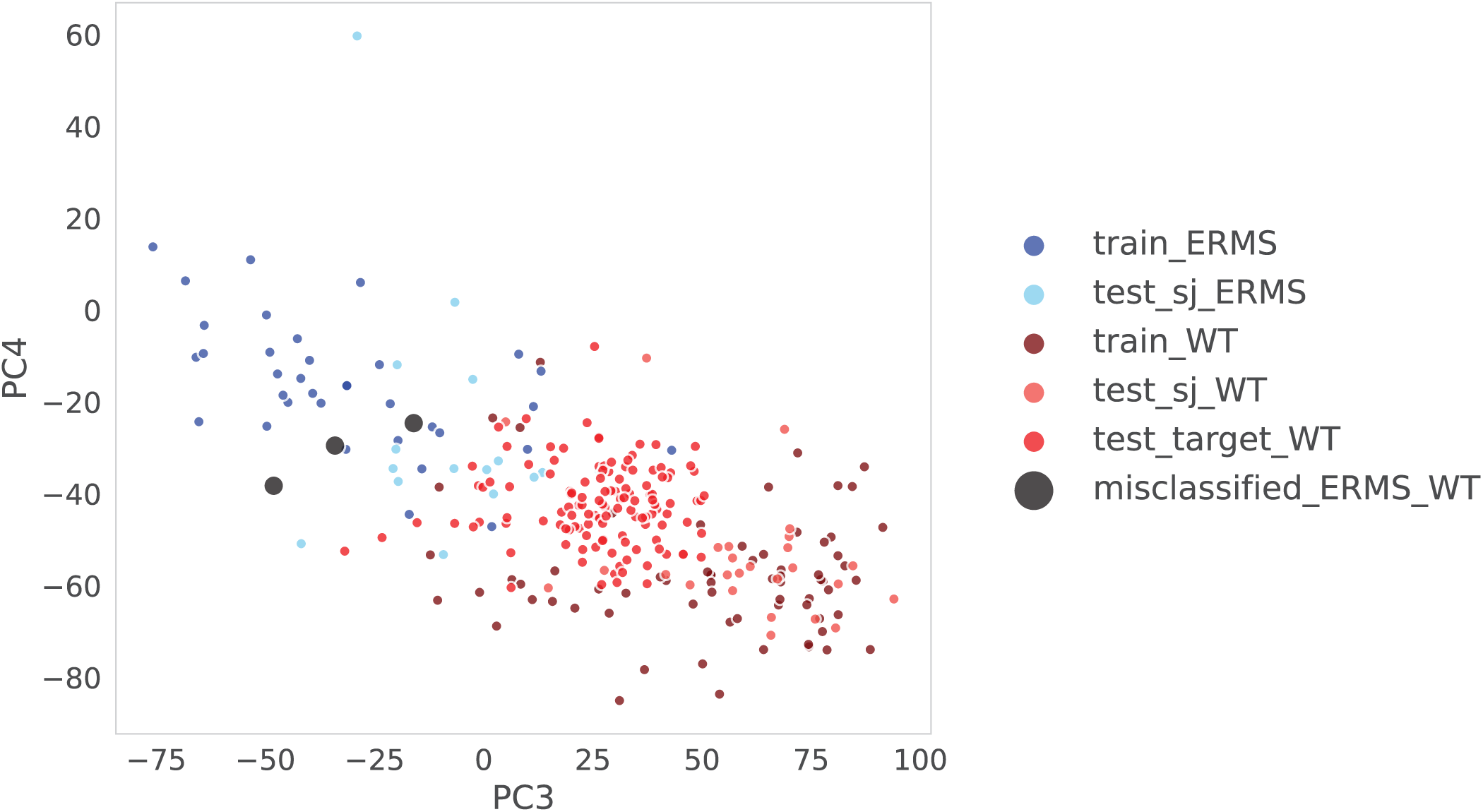
Principal component analysis highlights Wilms tumor cases misclassified as embryonal rhabdomyosarcoma. Samples shaded red denote WT, Wilms tumor; blue denotes ERMS, embryonal rhabdomyosarcoma. The three misclassified TARGET samples—CAAAAM, PAJMVC, and PAJNVX—labeled WT but predicted as ERMS, are shown as black points and cluster closer to the ERMS group than to WT.

**Supplementary Figure 12.**
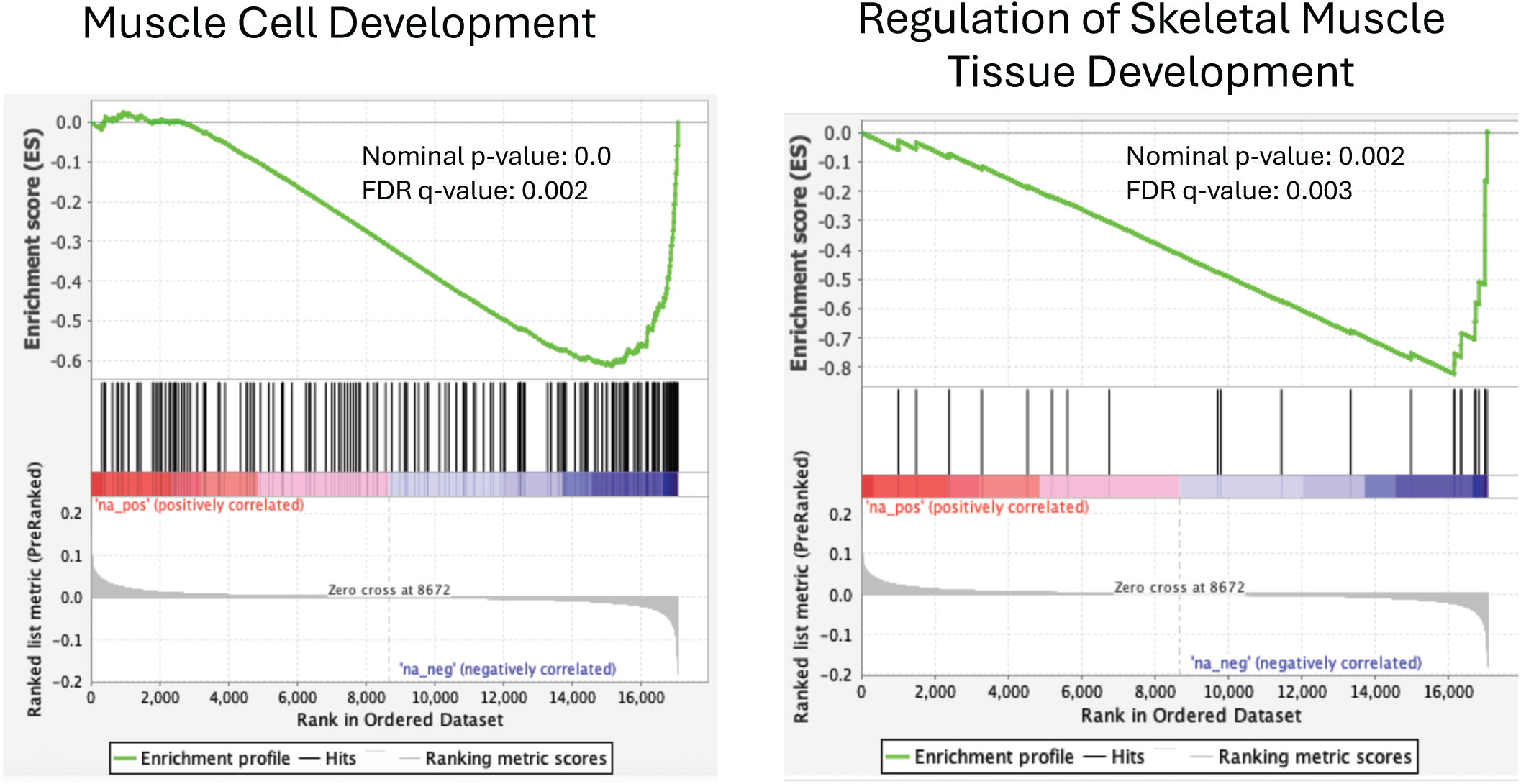
Principal component 3 captures myogenic differentiation programs. GSEA, gene set enrichment analysis show enrichment on the negative leading edge for pathways related to myogenic differentiation. This indicates that principal component 3 reflects variation driven by activation of muscle-lineage developmental programs in tumors.

**Supplementary Figure 13.**
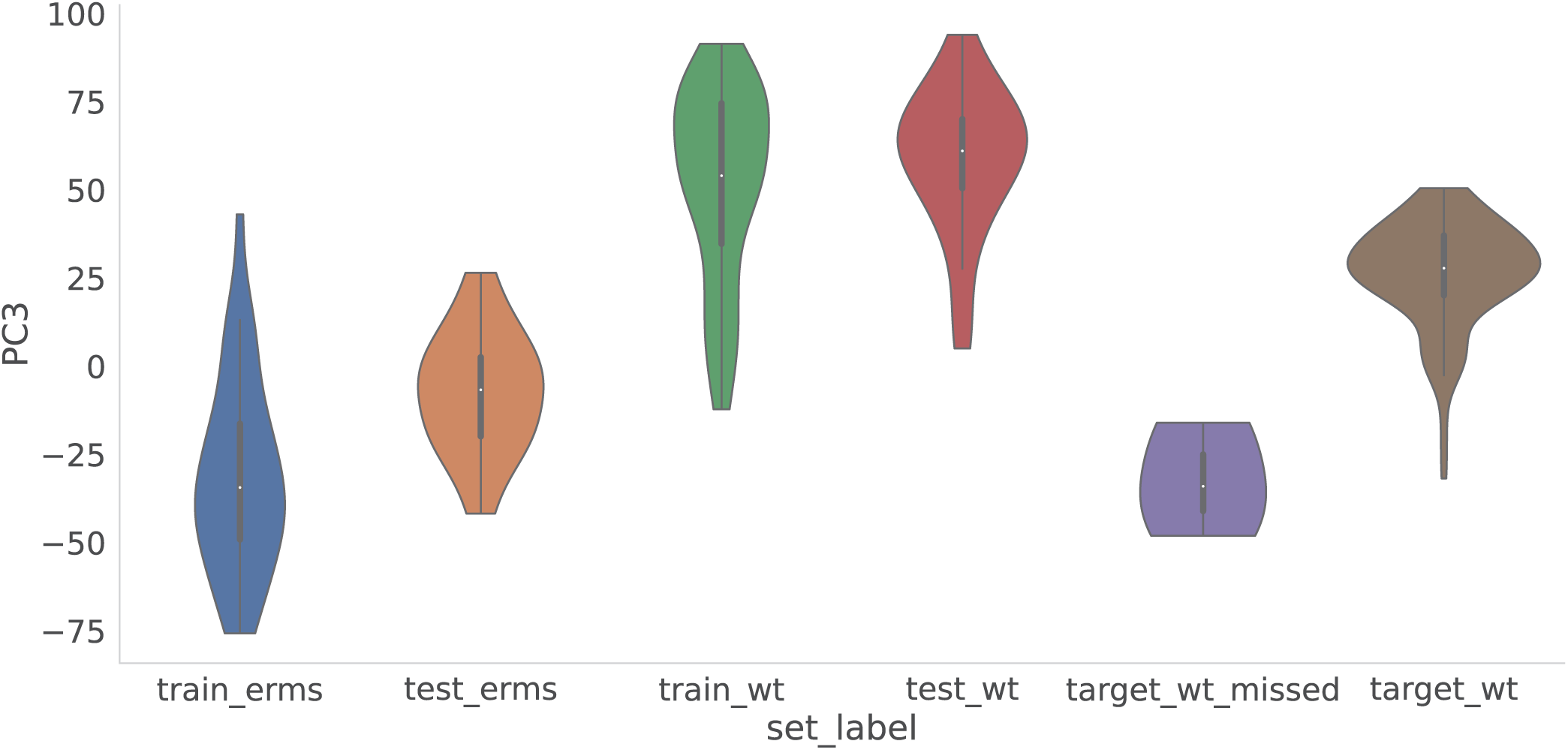
Principal component 3 distinguishes Wilms tumor from embryonal rhabdomyosarcoma. Violin plots of PC3, principal component values show that the three misclassified TARGET samples (CAAAAM, PAJMVC, PAJNVX) have negative PC3 values (−32.7 ± 13.1), aligning with embryonal rhabdomyosarcoma training samples (−24.0 ± 26.8) rather than Wilms tumor samples (38.6 ± 24.6).

**Supplementary Figure 14.**
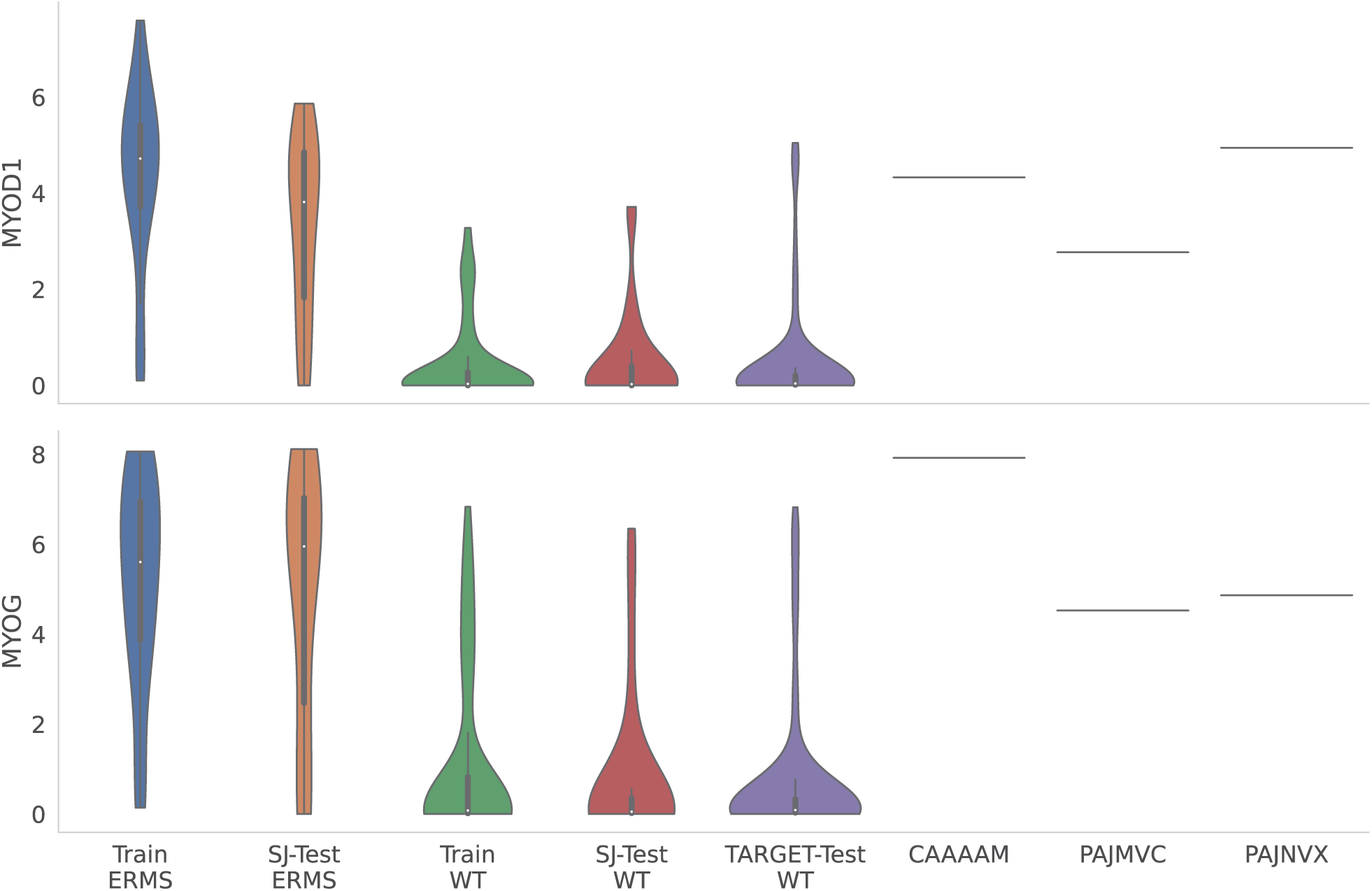
Elevated myogenic gene expression observed in Wilms tumor cases misclassified as embryonal rhabdomyosarcoma. Three Wilms tumor samples from the TARGET cohort—CAAAAM, PAJMVC, and PAJNVX—predicted as embryonal rhabdomyosarcoma show elevated *MYOD1* and *MYOG* expression, consistent with activation of myogenic differentiation programs. *MYOD1* and *MYOG* are master transcriptional regulators of myogenesis.

**Supplementary Figure 15.**
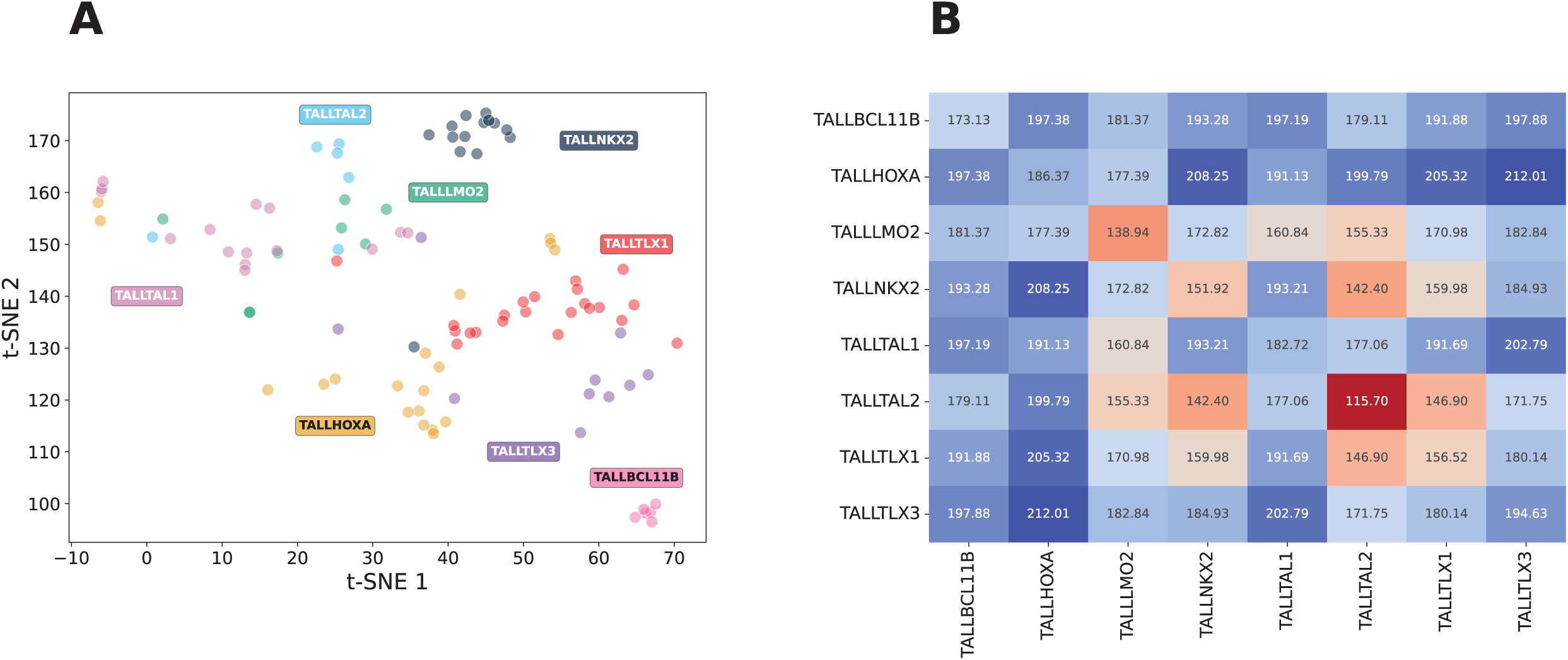
LMO2 T-cell acute lymphoblastic leukemia cases align with the TAL1 subtype. A t-SNE, t stochastic neighborhood embedding projection of training T-ALL samples derived from principal component features (A) shows TALL-LMO2 tumor classes co-cluster with TALL-TAL1 tumor classes rather than forming a distinct group. A heat map of mean pairwise Euclidean distances among training expression profiles (B) indicates TAL1 is closest to LMO2: the mean TAL1–LMO2 distance is lower than the TAL1–TAL1 within-subtype distance and lower than distances to all other T-ALL subtypes.

### Supplementary Tables

- **Supplementary Table 1.** Summary of the training, testing, and other groups used in CanID development.
- **Supplementary Table 2.** Distribution of tumor samples across cohort source.
- **Supplementary Table 3.** Classification performance of CanID across independent test datasets.
- **Supplementary Table 4.** Prediction outcomes for rare cases across cohorts.
- **Supplementary Table 5.** Summary of clinical and research projects contributing RNA-seq data, including sequencing protocol across solid tumor and hematologic malignancy datasets.
- **Supplementary Table 6.** ANOVA-based ranking of solid tumor principal components.
- **Supplementary Table 7.** Principal component feature counts required to explain variance in solid tumor and hematologic malignancy datasets.
- **Supplementary Table 8.** CanID Base Model Parameters of the ensemble classifier.
- **Supplementary Table 9.** Significant Principal Components in training data between mislabeled classes in TARGET Solid tumor samples.
- **Supplementary Table 10.** CanID out-of-bag prediction results for training samples.
- **Supplementary Table 11.** CanID prediction results for ambiguous or not otherwise specified cases from SJ-Cloud and TARGET cohorts.
- **Supplementary Table 12.** CanID and OTTER predictions for SJCloud rhabdomyosarcoma not otherwise specified cases, with pathologist review and fusion 2 status.
- **Supplementary Table 13.** CanID prediction results for low-confidence cases where pathologist review identified concerns regarding sample quality, limiting reliability of molecular subtyping.
- **Supplementary Table 14.** CanID results for rare subtypes, defined as diseases with fewer than 10 cases, across SJ-Cloud and TARGET cohorts.
- **Supplementary Table 15.** CanID predictions for TestData1, comprising approximately 30% of harmonized SJ-Cloud patient samples.
- **Supplementary Table 16.** CanID prediction results for TestData2, an independent external validation set from TARGET.
- **Supplementary Table 17.** CanID prediction results for TestData3, Clinical Pilot samples, with comparison to OTTER predictions.
- **Supplementary Table 18.** CanID prediction results for rare Clinical Pilot cases.
- **Supplementary Table 19.** Gene set enrichment analysis of the solid tumor gene loadings for principal component 2.
- **Supplementary Table 20.** Gene set enrichment analysis of the solid tumor gene loadings for principal component 1.
- **Supplementary Table 21.** CanID prediction results, with comparisons across alternative alignment and genome annotation strategies.
- **Supplementary Table 22.** CanID prediction results compared across alternative alignment and genome annotation strategies.
- **Supplementary Table 23.** Gene set enrichment analysis of the solid tumor gene loadings for principal component 3.
- **Supplementary Table 24.** CanID classification accuracy across tumor subtypes represented by both mRNA capture protocols: polyA-derived and total RNA-derived samples.

